# Single-embryo phosphoproteomics reveals the importance of intrinsic disorder in cell cycle dynamics

**DOI:** 10.1101/2021.08.29.458076

**Authors:** Juan M Valverde, Geronimo Dubra, Henk van den Toorn, Guido van Mierlo, Michiel Vermeulen, Albert J.R. Heck, Puck Knipscheer, Liliana Krasinska, Daniel Fisher, Maarten Altelaar

**Affiliations:** Biomolecular Mass Spectrometry and Proteomics, Bijvoet Center for Biomolecular Research and Utrecht Institute for Pharmaceutical Sciences, University of Utrecht, Utrecht, 3584 CH Utrecht, Netherlands; Netherlands Proteomics Center, Padualaan 8, 3584 CH Utrecht, Netherlands; IGMM, University of Montpellier, CNRS, Inserm, Montpellier, France; Equipe Labellisée LIGUE 2018, Ligue Nationale Contre le Cancer, Paris, France; Department of Molecular Biology, Faculty of Science, Radboud Institute for Molecular Life Sciences, Oncode Institute, Radboud University Nijmegen, 6525 GA Nijmegen, the Netherlands; Oncode Institute, Hubrecht Institute–KNAW and University Medical Center, Utrecht, 3584 CT, Netherlands

**Author notes:** Equal contributions (Juan M Valverde, Geronimo Dubra, Liliana Krasinska, Daniel Fisher, Maarten Altelaar).

## Abstract

Switch-like cyclin-dependent kinase (CDK)-1 activation is thought to underlie the abruptness of mitotic onset, but how CDKs can simultaneously phosphorylate many diverse substrates is unknown, and direct evidence for such phosphorylation dynamics *in vivo* is lacking. Here, we analysed protein phosphorylation states in single *Xenopus* embryos throughout synchronous cell cycles. Over a thousand phosphosites were dynamic *in vivo*, and assignment of cell cycle phases using egg extracts revealed hundreds of S-phase phosphorylations. Targeted phosphoproteomics in single embryos showed switch-like mitotic phosphorylation of diverse protein complexes. The majority of cell cycle-regulated phosphosites occurred in CDK consensus motifs, and 72% located to intrinsically disordered regions. Dynamically phosphorylated proteins, and documented substrates of cell cycle kinases, are significantly more disordered than phosphoproteins in general. Furthermore, 30-50% are components of membraneless organelles. Our results suggest that phosphorylation of intrinsically disordered proteins by cell cycle kinases, particularly CDKs, allows switch-like mitotic cellular reorganisation.

## Introduction

Eukaryotic cell cycle progression depends on the CDK1-subfamily of CDKs and is presumed to arise from the collective behaviour of altered protein phosphorylation states. With the notable exception of CDK1, most CDK and cyclin genes are dispensable for cell proliferation in the majority of cell types in the mouse (Liu et al., 2017; Santamaria et al., 2007), while in fission yeast, oscillating activity of CDK1 alone can drive the entire cell cycle (Coudreuse and Nurse, 2010; Fisher and Nurse, 1996). This suggests that a rather limited core network of CDKs can drive the eukaryotic cell cycle, and that changes in overall CDK activity somehow determine the sequence of the complex processes required to duplicate the genome and distribute cellular components during cell division. This “quantitative model” (Fisher and Nurse, 1996), implies that there exist low and high overall CDK activity thresholds for entry into S-phase and mitosis, respectively, determined by the CDK-regulatory network. This network involves positive and double-negative feedback loops, as well as futile cycles of CDK and CDK-opposing phosphatase activity (Fisher et al., 2012). Mathematical modelling shows that such features of network organisation can generate ultrasensitivity and hysteresis in CDK1 activation (Novak et al., 2010), while the resulting bistability of CDK1 activity leads to a switch-like G2/M transition (Tyson and Novak, 2001). These theoretical concepts are supported by experimental evidence in *Xenopus* egg extracts and mammalian cells (Rata et al., 2018; Novak et al., 2010; Trunnell et al., 2010; Kim and Ferrell, 2007; Pomerening et al., 2003; Sha et al., 2003).

The presumed switch-like dynamics of the CDK1 regulatory network is consistent with the abrupt morphological reorganisation of the cell at mitosis. In metazoans, the nuclear envelope and lamina breaks down and many cellular structures are rapidly disassembled. These include nuclear pore complexes, nucleoli, pericentriolar material, splicing speckles, Cajal bodies, promyelocytic leukaemia (PML)-nuclear bodies and stress granules (Banani et al., 2017; Hyman et al., 2014; Shin and Brangwynne, 2017; Woodruff et al., 2018), which have been collectively referred to as membraneless organelles (MLO). Thus, MLO assembly and disassembly occurs in a cell cycle-dependent manner. MLOs are thought to assemble by mechanisms involving multivalent interactions between intrinsically-disordered regions (IDR) of proteins (Banani et al., 2017), and this process can be regulated by protein kinases, including CDKs (Berchtold et al., 2018; Hur et al., 2020; Rai et al., 2018; Yahya et al., 2021). Protein phosphorylation in general is enriched in IDRs (Iakoucheva et al., 2004) and this also appears to be true for CDKs (Holt et al., 2009; Michowski et al., 2020; Moses et al., 2007). As such, an attractive model is that CDK-mediated IDR phosphorylation might trigger rapid dissolution of many MLOs at mitosis. This would be consistent with the fact that CDK1-family CDKs can phosphorylate hundreds of sites on diverse proteins (Blethrow et al., 2008; Chi et al., 2008; Errico et al., 2010; Ubersax et al., 2003), and regulate DNA replication, mitosis, transcription, chromatin remodeling, DNA repair, the cytoskeleton, nuclear transport, protein translation, formation of a mitotic spindle and even ciliogenesis (Hydbring et al., 2016; Krasinska and Fisher, 2018; Lim and Kaldis, 2013).

Direct evidence for switch-like dynamics of cell cycle-regulated phosphorylation *in vivo* is currently lacking. Single-cell proteomics studies (Budnik et al., 2018; Lombard-Banek et al., 2019) have insufficient sensitivity and reproducibility for low stoichiometry and highly dynamic targets such as phosphosites. Therefore, studies analysing cell cycle phosphorylation have generally used cells blocked at different stages of the cell cycle to generate “snapshots” of the phosphorylation landscape (Olsen et al., 2010). However, highly dynamic phosphorylation states cannot readily be determined from populations of cells (Purvis and Lahav, 2013). Moreover, whole-culture synchronisation methods generate artefacts due to cell cycle perturbation (Cooper, 2019; Ly et al., 2015). This might explain why, in an *in vivo* phosphoproteomics study in fission yeast synchronised by chemical block and release of CDK1, overall cell cycle phosphorylation dynamics appeared progressive rather than switch-like (Swaffer et al., 2016). Alternative phosphoproteomics approaches on unsynchronised cells selected with centrifugal elutriation (Ly et al., 2014) or FACS (Ly et al., 2017), lack the temporal resolution to determine the dynamics of protein phosphorylation throughout the cell cycle.

Here, we overcame these obstacles by using an extremely sensitive phosphopeptide enrichment strategy (Post et al., 2017) to perform quantitative phosphoproteomics on the highly synchronous early cell cycles of *Xenopus laevis* embryos, which consist solely of S and M-phase (Newport and Kirschner, 1982, 1984). By performing parallel phosphoproteomics using synchronously replicating or mitotic egg extracts we could attribute cell cycle behaviour of individual sites. This allowed us to investigate the general features of cell cycle-regulated phosphorylation compared to the entire phosphoproteome, revealing the importance of intrinsic disorder. We next compiled high-confidence CDK substrates in human and yeast, and analysed disorder on a proteome-wide scale in all three species. Our data provide evidence for switch-like mitotic phosphorylation of multiple subunits of protein complexes involved in diverse biological processes, and suggest that CDKs control these dynamics by phosphorylating IDRs, which constitute a large proportion of cell cycle-regulated phosphorylations. The tools and resources we have developed should be useful in understanding the conserved principles of eukaryotic cell cycle control.

## Results

### In vivo *phosphoproteomics reveals dynamics of cell cycle-regulated phosphorylation*

To investigate cell cycle-regulated phosphorylation in an unperturbed *in vivo* system, we analysed individual *Xenopus laevis* embryos undergoing highly synchronous cell cycles of early development. We collected single embryos at 18 time-points separated by 15-minute intervals, while recording visual cues of cortical rotation of fertilised eggs and subsequent cell divisions. Phosphopeptides from each embryo were purified, separated by nano-LC and analysed by high-resolution mass spectrometry (Fig. 1A). This identified 4583 phosphosites with high localisation probability (>0.75) mapping to 1843 proteins (Fig. 1B; Data S1), the majority being phosphoserines (Fig. 1C). Individual embryo phosphorylation states strongly correlated, demonstrating their synchrony and the robustness of our methodology (Fig. 1D). We thus generated a cell cycle map of protein phosphorylation from an unfertilised egg to a 16-cell embryo.

**Figure 1.**
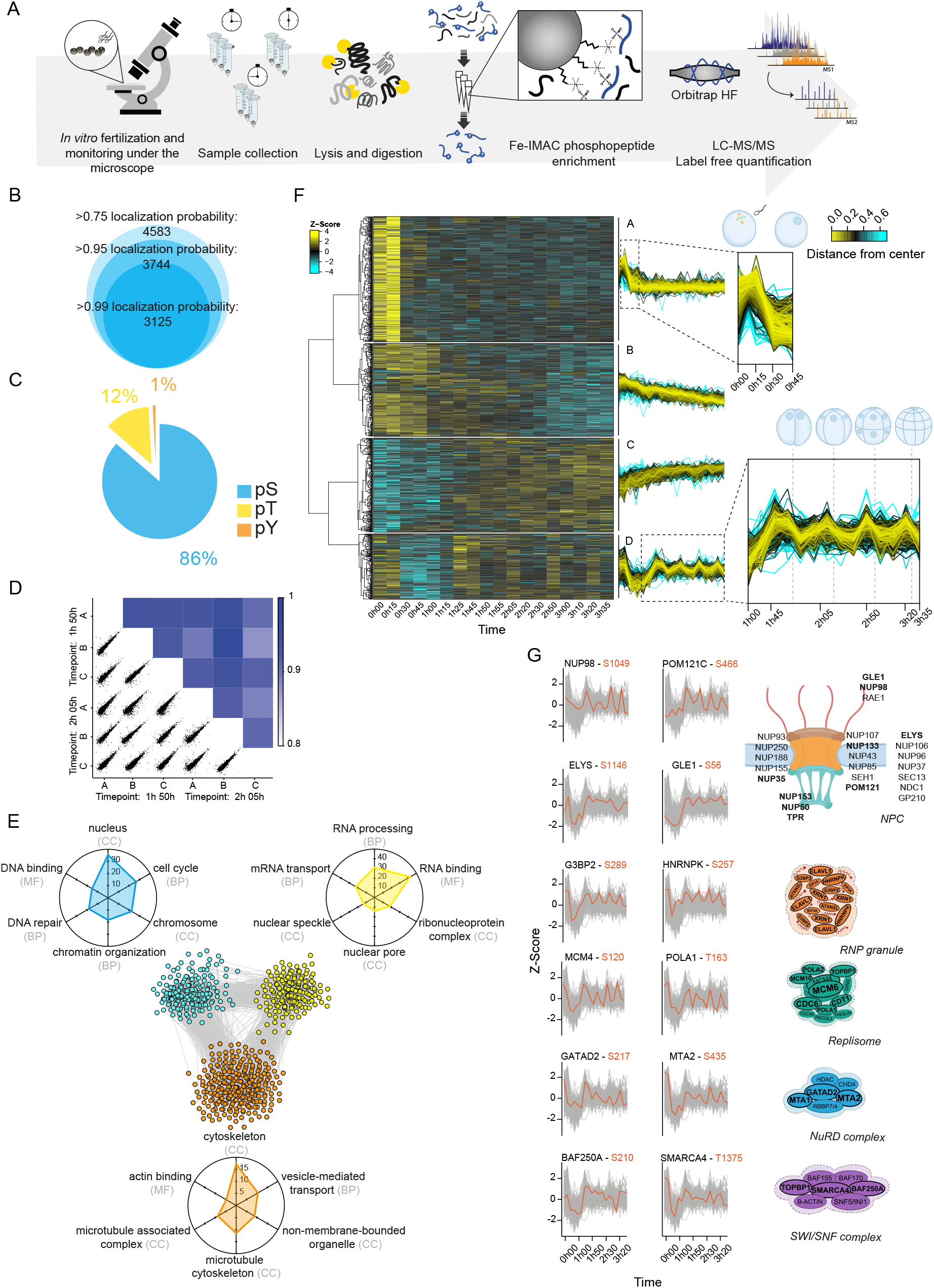
The time-resolved phosphoproteome from a single-cell to a 16-cell embryo. (A) Schematic representation of the workflow. Single *Xenopus* eggs and embryos were collected followed by cell lysis, protein digestion, phosphopeptide enrichment and high-resolution proteomics analysis. (B) Total number of phosphosites detected and their distribution according to the site localisation probability score. (C) Distribution of phosphosites identified among serine, threonine and tyrosine residues. (D) Correlation coefficients for two randomly selected time points. (E) STRING network of functionally associated proteins undergoing dynamic phosphorylation (each node represents a protein). Vicinity clustering reveals three main groups (yellow, blue and orange) with a high degree of association. Radar plots show the corresponding GO terms (adjusted p value <0.05) for each group (axes show -Log10(adj p value) for each GO term). (F) Hierarchical clustering of significantly changing phosphosites (ANOVA, Benjamini-Hochberg correction, FDR 0.05), reveals 4 clusters with distinct regulation (A-D). Dashed boxes in clusters A and D are zoomed-in to highlight dynamic phosphorylation patterns (dashed lines depict the time points of cell division). (G) Examples of proteins with known association showing similar oscillating phosphorylation. Plots highlight the dynamic trend of the cluster (grey) and selected phosphosites (orange) over time. Right, illustrations of protein complexes formed by the proteins undergoing dynamic phosphorylation. Proteins highlighted in bold show at least one oscillating phosphosite in our dataset.

We focused on 1032 sites on 646 proteins whose variation in abundance over time was statistically significant (ANOVA, Benjamini-Hochberg correction, FDR 0.05). Gene ontology (GO) and network analysis revealed high functional association and interconnectivity between groups of proteins involved in RNA binding and the nuclear pore complex (NPC), DNA replication and chromatin remodeling, and microtubule regulation (Fig. 1E). Hierarchical clustering of dynamic behaviour of the sites, in the absence of information on cell cycle timing, was sufficient to reveal four distinct groups that appear to reflect cell cycle-regulated behaviour (Fig. 1F; Data S1). Cluster A contained phosphosites with initial high intensity that dropped at 30 minutes, correlating with the degradation of cyclin B around 15 minutes after fertilisation, inactivation of the Mos/MAP kinase pathway, and exit from meiotic metaphase II. GO analysis for group A highlighted proteins involved in RNA regulation and nuclear organisation, including the NPC complex and nuclear transport, chromosomal structure and segregation (Fig. S1), as also observed in a recent study on meiosis exit in *Xenopus* eggs (Presler et al., 2017). Cluster B phosphosites were of lower intensity and dephosphorylation rate, and were enriched in regulators of RNA biosynthesis and stability, translation, actin, DNA replication and repair (Fig. S1). The levels of cluster A and B phosphosites were highest in eggs and just after fertilisation, and decreased during the first round of DNA replication, coincident with cortical rotation and transport of maternal mRNA responsible for axis specification in the embryo. This behaviour reflects the transition from meiosis to mitosis and suggests that dephosphorylation of these sites may prepare the zygote for upcoming cell divisions (Clift and Schuh, 2013).

Cluster C phosphosites progressively increased after meiotic exit, showing minor variations over the time course, while cluster D phosphosites had an oscillating signature with a clear upregulation preceding each cell division. GO analysis of cluster C reveals dominance of interphase cell cycle processes including DNA replication, RNA-related processes and chromosome organisation (Fig. S1). Cluster D was highly enriched in chromosome organisation and segregation, DNA replication, mRNA regulation, translation and microtubule binding. Cluster C included phosphosites displaying a reciprocal oscillating trend and a lower amplitude compared to cluster D sites, suggesting that they are phosphorylated during S-phase. Several sites with this trend, for example S31 of the replication licensing protein MCM4, were from monophosphorylated peptides, while the corresponding multiphosphorylated forms were found in cluster D (Fig. S1B). This suggests that cluster C contains the earliest phosphorylations of proteins that are highly phosphorylated towards mitosis. In cluster D, which peaks just before cell division and is enriched in proteins with roles in mitosis (Fig. S1), coordinated phosphorylation of multiple members of protein complexes involved in diverse processes commonly occurred, suggesting a common mechanism of regulation (Fig. 1G). Importantly, phosphoproteome changes were not simply a reflection of changes in abundance of the corresponding proteins (Fig. S2), which are comparatively negligible during *Xenopus* early development (Peuchen et al., 2017).

### High resolution phosphoproteomics reveals switch-like mitotic phosphorylation in vivo

The above results suggest that unsupervised clustering of phosphosite behaviour in single *Xenopus* embryos allows the resolution to detect mitotic sites. We investigated mitotic phosphorylation at high temporal resolution. A widely-held view is that mitotic phosphorylation is highly ordered due to specificity in CDK-substrate interactions (Örd et al., 2019), yet theoretical modelling suggests that it should occur in a switch-like manner due to the bistable mitotic CDK control network (Krasinska et al., 2011). Previous experimental data for CDK-dependent phosphorylations in synchronised cells shows a rather progressive increase throughout S-phase and G2 (Swaffer et al., 2016), but we suspected that this may be due to incomplete cell synchronisation (Ly et al., 2017). To see whether mitotic phosphorylation of individual phosphosites is progressive or switch-like *in vivo*, we analysed dynamics of cluster D sites in single embryos every 180-seconds using quantitative targeted phosphoproteomics (Lawrence et al., 2016; Schmidlin et al., 2016, 2019) by parallel reaction monitoring (Peterson et al., 2012). We thus obtained an extremely high-time resolution quantitative description of mitotic phosphorylation *in vivo* (Fig. 2A). We measured 64 phosphosites on proteins present in RNP granules, the replisome, chromatin remodeling complexes and NPCs. For each, we used heavy isotope-labeled phosphopeptides as internal standards. The results revealed parallel and abrupt upregulation of all phosphosites preceding each cell division (Fig. 2B, C), displaying switch-like phosphorylation of these diverse protein complexes at mitotic onset. These mitotic oscillations occurred despite global downregulation of CDK1 tyrosine-15 inhibitory phosphorylation (Fig. 2D), which agrees with biochemical evidence for downregulation of this phosphosite during early embryogenesis (Tsai et al., 2014). Analysis of the global embryonic phosphoproteome revealed an upward trend for phosphorylation of CDC25A during the first cell cycles (Fig. S3A), while other CDC25 homologues did not show statistically significant variations (CDC25B) or had no detected phosphosites (CDC25C). Phosphorylation of the maternally-expressed WEE1 homologue, Wee1-like protein kinase 2-A (hereafter WEE1A), varied only slightly after the first cell cycle (Fig. S3A). In contrast, there was a strong oscillation of phosphorylations on ubiquitin E3-ligases that control mitotic cyclin accumulation – NIPA, and the APC/C subunit APC1 – as well as Greatwall kinase, which regulates the PP2A inhibitors Arpp19/ENSA (Fig. S3B). Taken together with previous reports (Krasinska et al., 2011; Rata et al., 2018; Kamenz et al., 2021), these data suggest that control of mitotic cyclin levels and PP2A activity suffices for switch-like mitotic phosphorylation whereas regulated CDK1Y15 phosphorylation is not essential.

**Figure 2.**
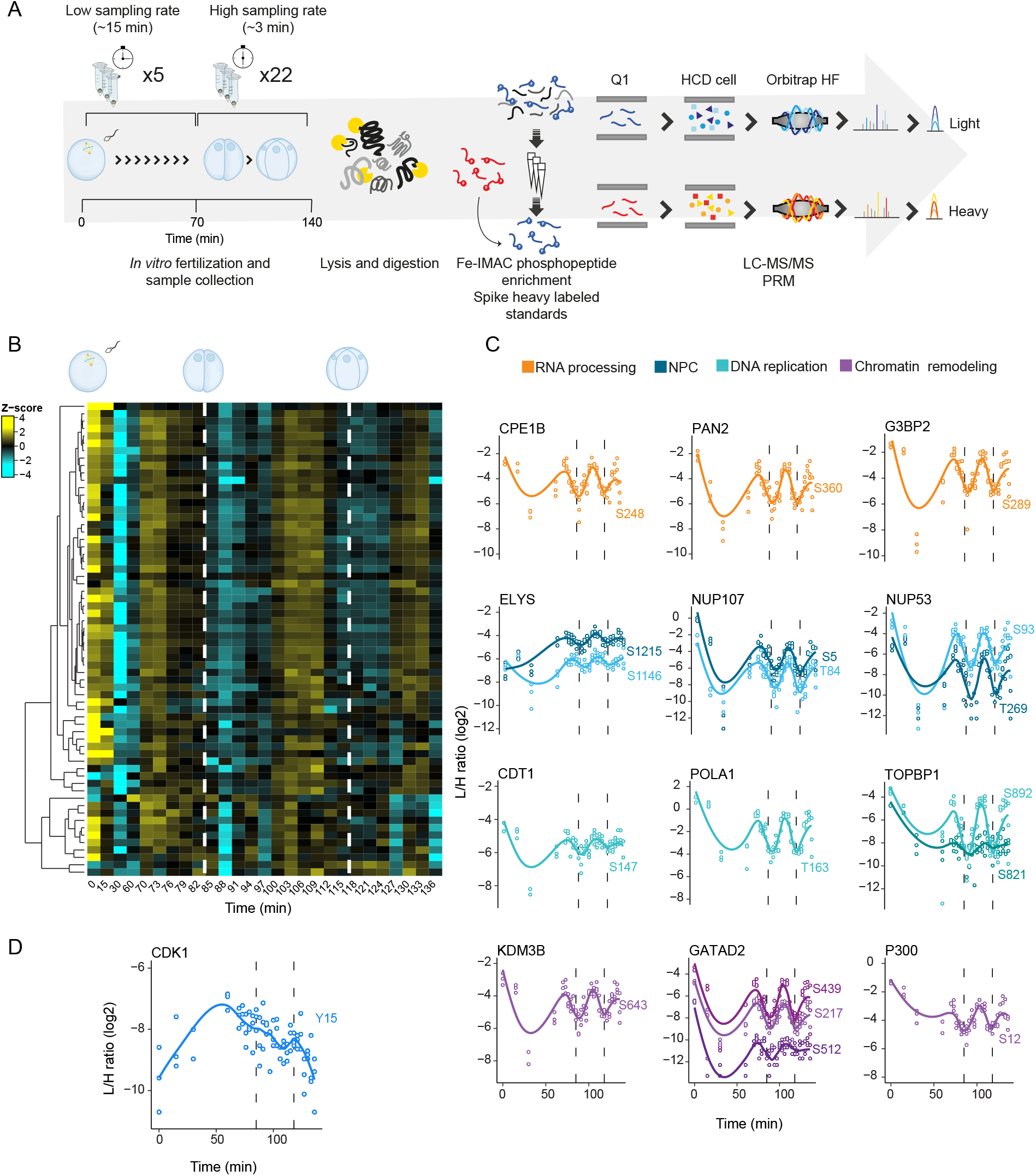
High-resolution targeted phosphoproteomics reveals switch-like mitotic phosphorylation *in vivo*. (A) Schematic representation of the workflow. Samples were collected over two cell divisions and enriched phosphopeptides were subjected to targeted proteomics analysis. (B) Heat map shows a highly synchronous wave of phosphorylation preceding each of the two cell divisions. Dashed lines depict times when cell divisions were recorded. (C) Single phosphosite plots from selected proteins. Each dot represents a biological replicate (n=3). Dashed lines depict times when cell divisions were recorded. (D) Single phosphosite plot of CDK1 inhibitory phosphorylation (Y15).

### Assigning phosphosites to different stages of the cell cycle

To assign embryo phosphosites to different cell cycle stages, we compared these *in vivo* phosphorylation patterns with protein phosphorylation states during a time course of DNA replication or in mitosis in egg extracts (Fig. 3A). Replication was initiated by adding purified sperm chromatin to interphase egg extracts and quantified (Fig 3B, top), while mitosis was triggered by adding recombinant cyclin B and verified microscopically. We also used egg extracts arrested at meiotic metaphase II (CytoStatic Factor, CSF, arrested). Overall, we identified 6937 phosphosites, which included 71% of the sites identified *in vivo* (Fig. 3C, Data S1). 1728 sites varied between replication and M-phase, including 693 sites upregulated in S-phase and 1035 in mitosis (Fig. 3B, Data S1). GO analysis of interphase sites revealed proteins involved in DNA replication, nuclear pore function, RNA polymerase binding, G2/M transition, chromosome and centromere organisation, cytoskeleton, and intracellular non-membrane-bound organelles (Fig. S4A). This phosphoproteomics dataset greatly increases the known repertoire of phosphorylation sites changing during S-phase.

**Figure 3.**
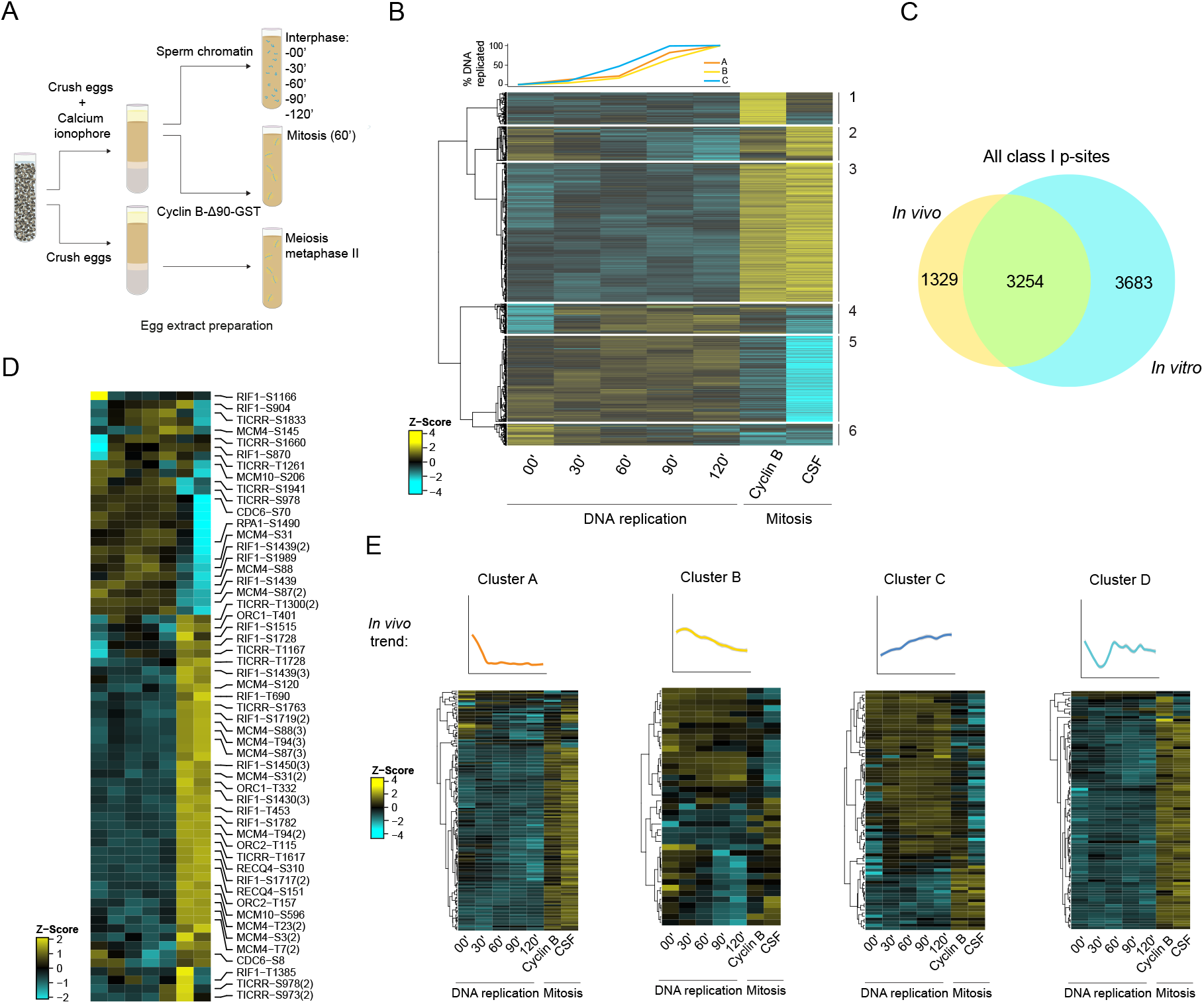
Phosphoproteome dynamics during DNA replication and mitosis *in vitro*. (A) Scheme of the experiment. (B) Top: quantification of DNA replication in each biological replicate. Below: Hierarchical clustering of dynamic phosphosites (ANOVA, Benjamini-Hochberg correction, FDR 0.05) reveals differential regulation of phosphosites during S-phase and mitosis. (C) Overlap between *in vivo* (embryo) and *in vitro* (egg extract) phosphoproteomics. (D) Heatmap of dynamic phosphosites detected in DNA replication factors. (E) Behaviour of *in vivo* dynamic phosphosites (top) in *in vitro* experiments.

We focused on the behaviour of phosphosites on proteins whose phosphorylation was previously described to promote DNA replication in vertebrates (Fig. 3D). We identified several sites on RECQ4 and Treslin/TICRR, ORC1 and ORC2 subunits of the origin-recognition complex, CDC6 (the catalytic subunit of DNA polymerase alpha), MCM10 and the single-stranded DNA binding protein, RPA1. We also detected 2 sites on MTBP, the metazoan homologue of yeast Sld7 (Kumagai and Dunphy, 2017), whose phosphorylation is required for DNA replication in a purified system (Yeeles et al., 2015), but these did not vary significantly throughout the time-course. The majority of replication proteins were phosphorylated on 1-3 sites during S-phase, with most phosphosites upregulated in mitosis. Only a few proteins were highly phosphorylated: on the MCM2-7 helicase, 21 sites were dynamic, most of which occurred on a single subunit, MCM4, while 17 dynamic sites were also present on RIF1, which opposes DDK-mediated MCM phosphorylation (Alver et al., 2017).

We next analysed the cell cycle behaviour of dynamic phosphosites that we found *in vivo* (Fig. 3E). Most embryo cluster A sites were upregulated in both CSF-arrested meiotic extracts and mitotic extracts, highlighting the functional similarities between meiotic and mitotic M-phase. There were few mitosis-specific sites but a significant fraction of sites that were specific for meiosis, reflecting the additional kinases active in meiotic M-phase, such as those of the Mos/MEK/MAP kinase pathway. Around half of embryo cluster B sites were present in interphase and absent in mitosis, while the rest showed a minimum phosphorylation in late S-phase, confirming that cluster B phosphosites are related to exit from meiotic metaphase II and are dephosphorylated during the first round of DNA replication. As expected, the majority of sites from embryo clusters C and D were part of the *in vitro* S-phase and mitotic groups, respectively, showing that data from single embryos can successfully discriminate phosphorylation occurring during these two cell cycle phases. Furthermore, this *in vitro* dataset confirms that in mitosis, monophosphorylated species of some proteins are reduced (thus appear in embryo clusters B or C) because multisite phosphorylation emerges (Fig. S4B; see also Fig. S1B).

### Potential CDK sites dominate the cell cycle-regulated phosphoproteome

To identify probable kinases responsible for phosphorylations occurring in these experiments, we analysed kinase consensus motifs. Around 51% of all the detected phosphosites *in vivo* were proline-directed (S/T-P), thus, conform to the minimal consensus for CDK sites (Fig. 4A). This proportion increased to 60% among dynamic sites, with around 10% of all phosphosites matching the full canonical CDK1-family sequence motif S/TPxK/R. Phosphosites in replicating and mitotic extracts displayed a similar trend for minimal and full CDK consensus motifs (Fig. 4A). Putative CDK targets dominated all clusters, with over 80% of sites in cluster D *in vivo* and mitotic clusters *in vitro* conforming to at least the minimal CDK motif (Fig. 4B, Fig. S5A, B). While in meiotic M-phase, MAP kinases, which have the same consensus motif as CDKs, are likely responsible for a subset of these sites (*i*.*e*. those specific to embryo cluster A or CSF extracts), these kinases are inactivated upon meiotic exit and not reactivated during embryogenesis (Ferrell et al., 1991), suggesting that most of the dynamic proline-directed sites are due to CDKs.

**Figure 4.**
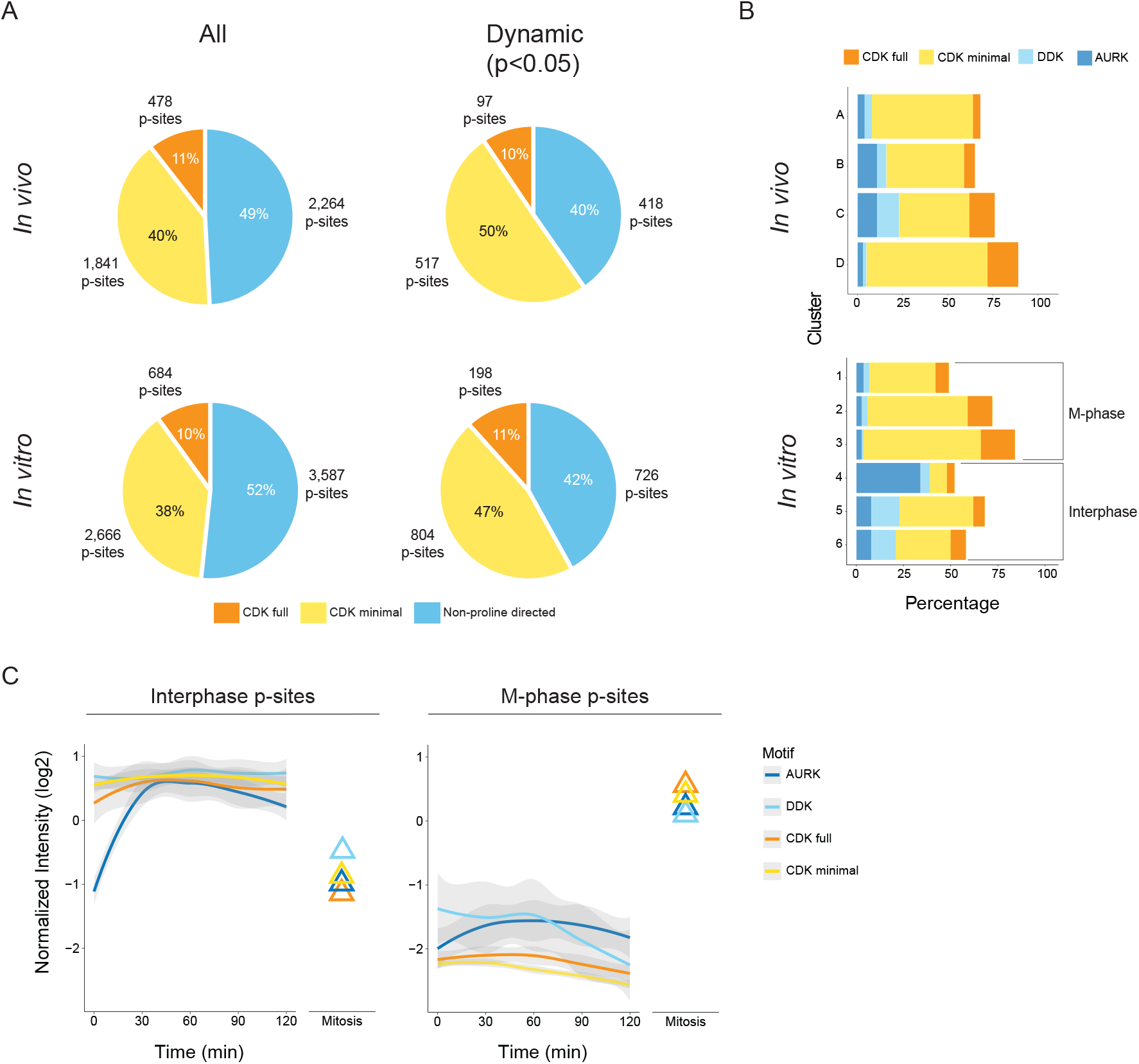
Proline-directed phosphosites dominate the early embryo phosphoproteome. (A) Distribution of potential CDK targets among all detected phosphosites and dynamic phosphosites, *in vivo* (embryo, top) and *in vitro* (egg extract, bottom). (B) Count of phosphosites according to their potential upstream kinase for each cluster in the *in vivo* (top) and *in vitro* (bottom) experiments. (C) Dynamic trend of phosphorylations of potential kinase targets in egg extract S-phase clusters (4-6, left) and in mitotic clusters (1-3, right).

Potential target sites of other kinases such as Aurora, PLK, DDK and Casein kinase I and II were also present, albeit to a lesser extent (Fig. S5A). Unexpectedly, consensus sites for Aurora were most enriched in cluster C *in vivo* and the interphase clusters *in vitro* (Fig. 4B; Fig S5), and these sites were upregulated early during the replication time course (Fig. 4C). This is consistent with kinetochore processes being enriched in the GO analysis, and suggests that many Aurora targets are phosphorylated prior to mitosis.

These results substantially increase our knowledge of vertebrate cell cycle-regulated phosphorylations, and show that, irrespective of the cell cycle phase, the most abundant phosphosites that change during the cell cycle are potential CDK targets. Taken together, our data suggest that CDKs are responsible for the majority of cell cycle-regulated phosphorylations.

### Xenopus *dynamic phosphoproteins and CDK substrates in human and yeast are highly disordered*

Although few direct CDK substrates have been characterised in *Xenopus*, we surmised that they are likely conserved between vertebrate species. We therefore compiled a set of 656 human CDK1-subfamily targets (Data S2), combining data on 450 CDK substrates from PhosphoSite Plus (Hornbeck et al., 2015) with manually curated information on 206 targets from several human CDK substrate screens and other studies (see Supplementary Methods for sources). 303 of these 656 CDK substrates have *Xenopus* homologues among the 1843 phosphoproteins we detected, and 149 were present among the 646 proteins with dynamic phosphosites in *Xenopus* embryos (Fig. 5A). Thus, the predominance of CDK motifs among dynamic phosphosites reflects a high proportion of *bona fide* CDK substrates. This may underestimate the true fraction, since we only considered proline-directed sites as CDK motifs, yet of the 1200 human CDK phosphosites on 656 substrates, 124 were non-proline-directed, confirming a previous report that CDK1 can also efficiently phosphorylate non-proline-directed sites (Suzuki et al., 2015). We estimated the likely fraction of the latter proteome-wide by taking advantage of the extensive data available for CDK1 targets in budding yeast, whose cell cycle regulation is conserved with vertebrates. We defined high-confidence CDK1 sites (Data S2) by intersecting data for *in vitro* CDK1 substrates (Ubersax et al., 2003) with *in vivo* CDK1-dependent phosphosites (Holt et al., 2009). 100 of the 185 yeast CDK1 substrates defined *in vitro* were also phosphorylated in a CDK1-dependent manner *in vivo*, but 19 of these were not proline-directed (Fig. S5C). Taken together with our data, this suggests that, as well as most proline-directed sites, a sub-fraction of dynamic non-proline-directed phosphosites in our dataset are mediated by CDKs, and reinforces the dominant role of CDKs in cell cycle-regulated phosphorylation.

**Fig. 5.**
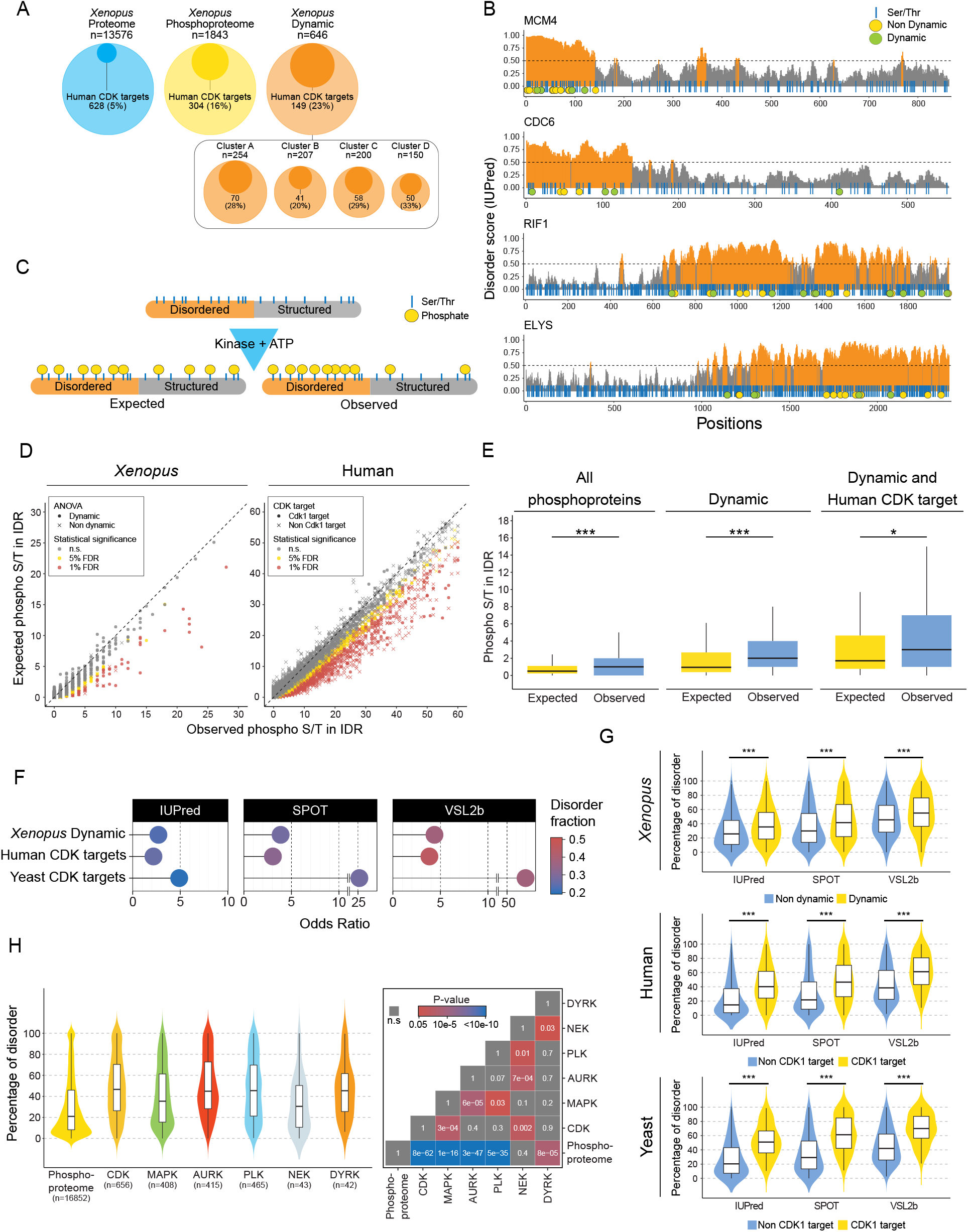
Dynamic phosphoproteins and CDK substrates are characterised by intrinsic disorder. (A) Circle plots presenting enrichment of homologues of human CDK substrates among *Xenopus* phosphoproteins detected *in vivo* and those with dynamic phosphosites. (B) Diagrams of IUPred scores over the length of selected proteins. Regions with scores >0.5 (orange) are considered to be disordered, and <0.5 (grey) structured. Blue vertical lines indicate Ser and Thr residues; yellow circles, phosphorylated sites; green circles, dynamic phosphosites. (C) Scheme illustrating hypothetical enrichment of phosphorylation in disordered regions when taking into account amino acid compositional bias. (D) Scatter plot of expected vs observed phosphorylated Ser/Thr for each protein of human and *Xenopus* phosphoprotein datasets. FDR thresholds of 5% and 1% are marked in yellow and red respectively. Crosses: proteins with at least one dynamic phosphorylation in *Xenopus*, or human CDK1 subfamily substrates, respectively. (E) Boxplots showing expected vs observed phosphorylated Ser/Thr among all phosphoproteins detected (left), phosphoproteins with at least one dynamic phosphosite (middle), and dynamic phosphoproteins also detected as CDK1 subfamily targets in humans (right). Distributions were compared with the Wilcoxon signed-rank test. *p<0.05, **p<0.01, ***p<0.001. (F) Plots showing the common Odds Ratio of Ser/Thr phosphorylation in structured and ordered regions calculated with the Fisher’s test (see Fig. S6B, C). For all organisms, the disordered regions were calculated with three different disorder predictors. The disordered fraction is presented in a colour scale. (G) Violin plots of the distribution of disordered residues per protein for CDK targets vs the rest of the phosphoproteome for human and yeast, and dynamic phosphoproteins vs the rest of the phosphoproteome for *Xenopus*. Intrinsic disorder was calculated with three different predictors (IUPred, SPOT, and VSL2b). Statistical significance was evaluated with the Wilcoxon–Mann–Whitney test. (H) Violin plot (left) showing the distribution of disordered residues per protein for CDK, MAPK, Aurora, PLK, NEK and DYRK kinase targets vs the rest of the phosphoproteome for human targets. Statistical significance was assessed by Krustal-Wallis ANOVA, and pairwise comparisons were performed with Dunn’s post-hoc tests. The adjusted p-values (Benjamini-Hochberg) are shown in a tile plot (right).

Next, we wondered whether the diverse dynamically phosphorylated proteins share common structural features facilitating CDK-mediated phosphorylation. Since phosphosites in general are often located in IDRs of proteins (Iakoucheva et al., 2004), we first computationally analysed intrinsic disorder in our *in vivo* dataset using the energy estimation-based predictor IUPred (Dosztanyi et al., 2005). We indeed found that phosphosites were often located in predicted IDRs, which was especially striking for highly phosphorylated proteins such as MCM4 and RIF1 (Fig. 5B). However, since sequence attributes of phosphorylation sites in general are similar to those found in IDRs (Iakoucheva et al., 2004), this finding, as well as previous observations of yeast and mouse CDK sites being preferentially located in IDR (Moses et al., 2007; Holt et al., 2009; Michowski et al., 2020), may at least partly result from the enrichment of serine, threonine and proline in disordered regions. To explore this possibility, we analysed intrinsic disorder among the entire phosphoproteome of *Xenopus*, human and yeast. To do this, we used three different prediction methods with differing sensitivity, including the conservative IUPred, a more permissive predictor, VSL2b (Peng et al., 2006), and an intermediate-sensitivity predictor, SPOT (Hanson et al., 2017). All three are widely used, and a recent systematic comparison ranked SPOT among the best-performing predictors (Necci et al., 2021). As we anticipated, all predictors found that phosphorylatable amino acids and proline are enriched in IDRs (Fig. S6A). To correct for this compositional bias and investigate intrinsic disorder among the cell cycle phosphoproteome systematically, we compared the number of phosphosites detected in predicted IDRs to that expected according to the distribution of phosphorylatable amino acids (Fig. 5C). Even after this correction, our identified phosphosites were strongly enriched in predicted IDRs, especially for proteins with at least one site displaying dynamic phosphorylation. The same was true for human CDK substrates (Fig. 5D, E). Therefore, CDKs more readily phosphorylate disordered regions of proteins.

To estimate the differential phosphorylation of disordered sites globally, we calculated the ratio of dynamically phosphorylated (*Xenopus*) or CDK-phosphorylated (yeast, human) to non-phosphorylated serine and threonine in both disordered and structured regions (Fig. S6B; see Methods). This ratio can be approximately equated with the relative probablity of a Ser/Thr to be phosphorylated when located in a disordered region compared with those in structured regions. For all predictors, this ratio was far higher than one for all species (Fig. 5F, Fig. S6C), confirming that phosphosite enrichment in IDRs is a general feature of cell cycle-regulated and CDK-mediated phosphorylations throughout eukaryotes. We then asked whether this is specific to CDKs or is also true for substrates of other major mitotic kinases. Using available data on their substrates from Phosphosite Plus and recent phosphoproteomics studies, we analysed the mitotic polo-like (PLK) and Aurora kinases. We also analysed substrates of the less well-known DYRK kinases, which are closely related to CDKs and have recently been found to promote mitotic phosphorylation of certain intrinsically disordered proteins (IDPs) (Rai et al., 2018), and the NEK kinase family, which has roles in centrosome duplication and various stages of mitosis. We compared sites of each mitotic kinase with sites of MAP kinases, which, although not generally considered cell cycle kinases, are evolutionarily related to CDKs and share the proline-directed S/T consensus site. For each kinase, the phosphosites were strongly enriched in IDR (Fig. S6D), supporting the idea that intrinsic disorder generally promotes phosphorylation (Iakoucheva et al., 2004).

To explain the dominance of CDK-mediated cell cycle phosphorylation, we surmised that proteins that are phosphorylated in a cell cycle-dependent manner, and CDK substrates in particular, might be more disordered overall than phosphoproteins in general. We therefore determined the percentage of disordered residues of *Xenopus* proteins with dynamic phosphosites, as well as human CDK substrates, compared to the rest of their respective phosphoproteomes (non-cell cycle-dependent phosphorylations in our *Xenopus* dataset, and non-CDK-dependent phosphoproteins from the Phosphosite Plus database) (Data S3). This revealed that, on average, both *Xenopus* dynamic phosphoproteins and human CDK substrates contain approximately twice as many disordered amino acids than the average of all other phosphoproteins (Fig. 5G), putting them among the top quartile of proteins with the most disorder in the proteome. High-confidence yeast CDK1 substrates are also highly enriched in disorder compared to the non-CDK-dependent yeast phosphoproteome (Fig. 5G), revealing conservation of this principle across eukaryotes. The same trend was observed using 13 different intrinsic disorder prediction methods, supporting the robustness of our conclusions (Fig. S6E). Almost all methods gave a lower fraction of disordered residues in the yeast proteome than in *Xenopus* and human (Fig. S6F), consistent with an increase in disorder in more complex organisms (Darling and Uversky, 2018). We wondered whether this exceptional level of disorder among CDK substrates compared to the overall phosphoproteome might reflect the importance of disordered proteins for the cell cycle in general. If so, then substrates of other cell cycle regulated kinases might also be enriched in disorder. We thus compared them with substrates of MAP kinase, whose phosphosites are also preferentially located in IDR (Fig. S6D), but which is generally involved in cell signaling rather than the cell cycle. This showed that, like CDK substrates, targets of the cell cycle kinases are highly enriched in disorder, and, with the exception of NEK substrates, are significantly more disordered than targets of MAP kinase (Fig. 5H). Relatively few NEK substrates are known, and these are the most structured substrates of the kinases analysed, but this conclusion might evolve as more NEK targets are identified.

### CDK targets are major constituents of MLOs

Protein phosphorylation regulates cell cycle-dependent assembly and disassembly of membrane-less organelles (MLO) (Berchtold et al., 2018; Hur et al., 2020; Rai et al., 2018). Since we observed clustering of cell cycle-regulated phosphosites in IDRs (Fig. 5B), we hypothesised that changes in the net charge of these regions due to mitotic phosphorylation might modulate liquid-liquid phase separation, which is thought to control MLO formation. To see whether the evidence supports this hypothesis, we first applied a recently-developed machine learning classifier (van Mierlo et al., 2021) to predict whether cell cycle-regulated phosphoproteins, or CDK substrates, have an increase in average propensity for phase separation (defined as PSAP score). Indeed, using *Xenopus* data, we observed a sharp increase in the PSAP score, from the proteome to the phosphoproteome, and a further increase for dynamic phosphoproteins, with the highest score for mitotic cluster D (Fig. 6A). We then similarly analysed human kinase substrates, which showed that the propensity for phase separation is far higher amongst targets of the cell cycle (CDK, Aurora, PLK, again, with the exception of NEK) and DYRK kinases than the overall phosphoproteome, but less so for MAP kinase substrates. We note that, unlike the *Xenopus* early embryo phosphoproteome, the human phosphoproteome did not show a striking difference in PSAP score to the proteome. This suggests that the early embryonic phosphoproteome is already highly enriched in proteins with a propensity to phase separate and in targets of cell cycle kinases.

**Fig. 6.**
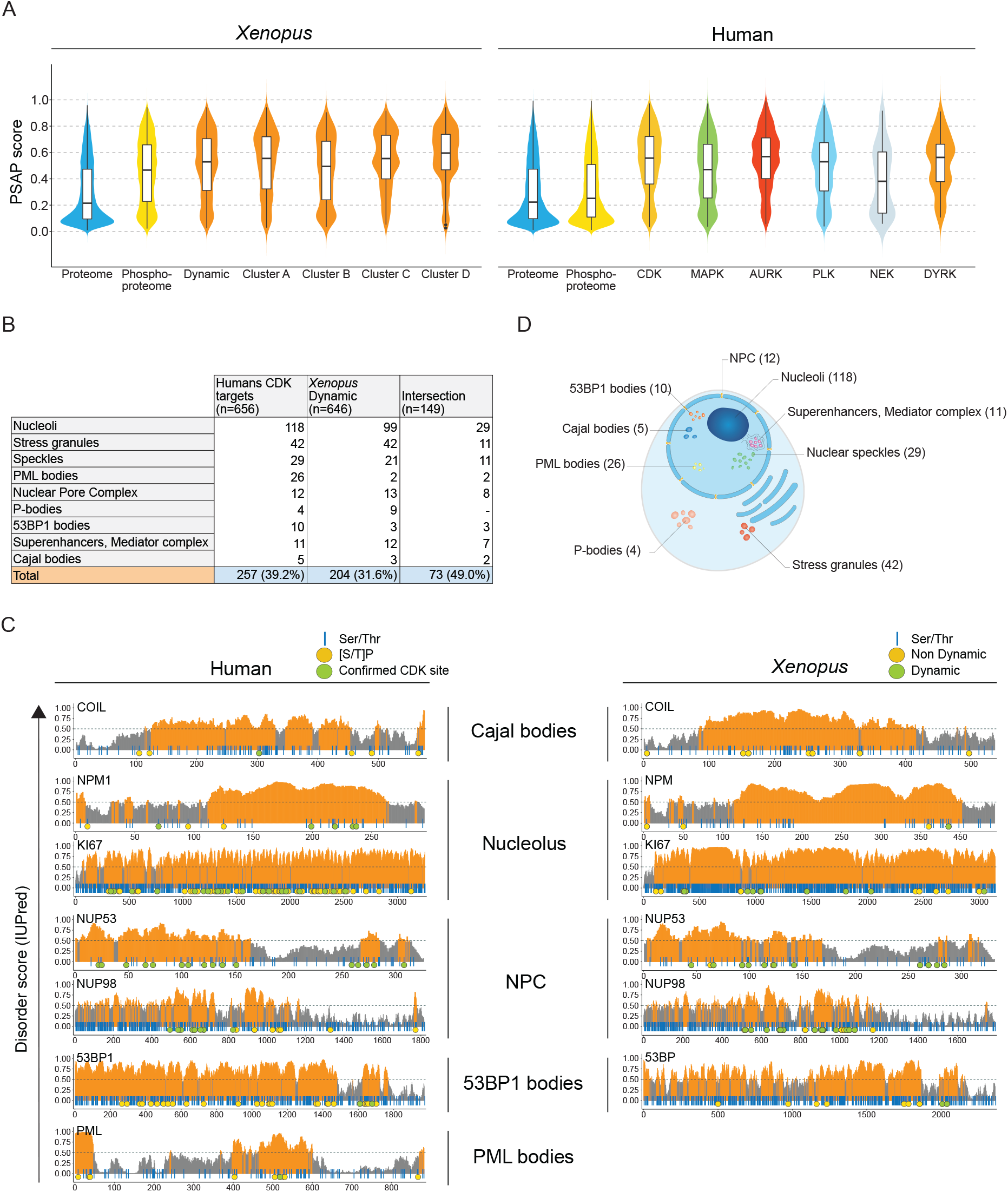
CDK targets are major MLO components. (A) Violin plots presenting PSAP score for *Xenopus* dynamic phosphoproteins (left), and human kinase targets, in comparison with total proteome and phosphoproteome. (B) Human CDK1 subfamily targets, *Xenopus* dynamic phosphoproteins, and the intersection of both sets, that are present in our manually curated proteome of membraneless organelles. (C) Diagrams of IUPred scores over the length of human CDK targets identified as primary components of MLOs in different studies, and their *Xenopus* homologues in this study. Regions with scores >0.5 (orange) are considered to be disordered, and <0.5 (grey) structured. Blue vertical lines indicate Ser and Thr residues; yellow circles, known Ser/Thr-Pro phosphosites (human) and non-dynamic phosphosites (*Xenopus*); green circles, confirmed CDK1 subfamily phosphorylations (human) and dynamic phosphorylations (*Xenopus*), from both embryos and egg extracts.

Next, we analysed publicly available data on each of our curated human CDK substrates for localisation to MLOs, concentrating on MLOs in which phase separation has been clearly demonstrated to play a role. We found that at least 257 CDK substrates (39.2%) are present in MLOs (Fig. 6B). We then assembled an MLO proteome from human proteomics studies (Data S4; See Supplementary Methods for sources), and analysed the proportion of *Xenopus* dynamic phosphoproteins with homologues among this dataset. This revealed that 204 dynamic *Xenopus* phosphoproteins (31.6%) are found in MLOs. CDK substrates include major proteins of MLOs highly enriched in IDPs such as coilin (Cajal bodies), nucleophosmin and Ki-67 (nucleoli), 53BP1 (53BP1 bodies), nucleoporins (NPCs) and PML (PML bodies). In these proteins, as in dynamic phosphoproteins *in vivo*, the vast majority of proline-directed phosphosites and confirmed CDK sites were located in predicted IDRs (Fig. 6C). Moreover, of the 149 proteins that show dynamic phosphorylation in *Xenopus* and are CDK substrates in human, 73 (50%) localise to MLOs (Fig. 6B). These data support the existence of an evolutionary conserved mechanism for cell-cycle control of MLOs.

## Discussion

In this work, we identify dynamics of cell cycle regulated phosphorylations in an unperturbed physiological system *in vivo* by applying a combination of single *Xenopus laevis* embryo phosphoproteomics at high temporal-resolution over synchronous cell divisions, and parallel phosphoproteomics in synchronously replicating or mitotic egg extracts. This unique approach reveals important insights into cell cycle regulation.

First, we find evidence that CDKs are responsible for the majority of cell cycle-regulated phosphorylations. We compiled a database of high-confidence human CDK1-family substrates and found that nearly a quarter of them are represented among the dynamically phosphorylated proteins in *Xenopus* embryos, while more than half the phosphosites conform to the minimal CDK consensus motif. This is likely to underestimate the true proportion since our human CDK substrate dataset is probably incomplete, and, like previous screens for CDK1-family substrates in different organisms (Ubersax et al., 2003; Moses et al., 2007; Swaffer et al., 2016; Michowski et al., 2020), we only considered proline-directed sites, yet CDK1 can also efficiently phosphorylate non proline-directed sites in certain contexts (Suzuki et al., 2015). We find that around 10% of human CDK phosphosites are non-proline-directed, while our analysis of high-confidence CDK substrates in yeast indicates that the corresponding figure may be closer to 20%. Even this may be a conservative estimate since it relied on analysis of phosphorylation of proteins that had been selected to contain several proline-directed consensus motifs. This illustrates a frequent bias in analysis of CDK sites, in that non-proline-directed sites are almost always filtered out.

Second, we demonstrate switch-like behaviour of mitotic phosphorylation *in vivo*, and its occurrence on entire protein complexes from a variety of biological processes. This is consistent with mathematical modelling of the mitotic CDK regulatory network, which predicts switch-like CDK1 activation. Surprisingly, however, this abrupt phosphorylation of many CDK substrates occurred despite progressive downregulation of CDK1 phosphorylation on tyrosine-15, which has been thought to be a key contributor to ultrasensitivity in CDK1 activation (Kim and Ferrell, 2007; Trunnell et al., 2010). Indeed, early work on cell cycles during *Xenopus laevis* development demonstrated that CDK1-Y15 phosphorylation was absent from cell cycles 2-12 (Ferrell et al., 1991), although more recent data suggest low level and minor fluctuations in cell cycles 2-3 (Tsai et al., 2014), consistent with our findings. We also observe similar fluctuations of S38 in Xe-Wee1A, and S120 and S299 of CDC25A, which are likely mediated by CDK1. Our data suggest that the CDC25 positive feedback loop is active and increasing over time, while the WEE1 double-negative loop remains constant, explaining the slight oscillation but progressive general downregulation of CDK1 Y15 in subsequent cell divisions.

If regulated CDK1-Y15 phosphorylation decreases over time, how then could the switch-like dynamics be sustained for a prolonged period after the first cell division? One possibility is that futile cycles of CDK1 and opposing protein phosphatase activity, likely PP2A (Krasinska et al., 2011), are responsible. The Goldbeter-Koshland model of futile cycling predicts switch-like changes in network output upon small variations in the relative activities of opposing enzymes around a critical threshold, even without any feedback between the opposing enzymes (Goldbeter and Koshland, 1981). PP2A is inhibited by Greatwall kinase-mediated phosphorylation of ARRP19/ENSA (Gharbi-Ayachi et al., 2010; Mochida et al., 2010), which is promoted by CDK1 itself (Yu et al., 2006). Consistent with our data and this model, reconstituting mitotic entry with purified components in *Xenopus* egg extracts demonstrated that switch-like behaviour did not depend on CDK1-Y15 regulation, but, rather, reciprocal regulation between CDK1 and PP2A via Greatwall (Mochida et al., 2016). A second mechanism which is independent of regulated CDK1-Y15 phosphorylation is positive feedback in cyclin B1 accumulation, since CDK1-mediated phosphorylation of the ubiquitin ligase subunit NIPA prevents interphase cyclin B1 degradation (Bassermann et al., 2005, 2007). Our data suggest that both CDK1-Y15-independent mechanisms may contribute to switch-like phosphorylation since we find that both Greatwall and NIPA phosphorylation oscillate at mitosis *in vivo*.

Third, we reveal phosphorylation dynamics of known DNA replication factors and many others during S-phase. MCM4 and RIF1, which are involved in DDK-mediated initiation of DNA replication and replisome stability (Alver et al., 2017), are among those with the highest number of regulated phosphosites, suggesting a role for hyperphosphorylation of a subset of replication proteins. As expected, we find that potential DDK phosphosites are upregulated during S-phase, both *in vivo* and in *in vitro*. We observe a similar trend for putative targets of Aurora kinases, best known as regulators of the microtubule-kinetochore attachment at prometaphase and of cytokinesis. This suggests that phosphorylation of many of their substrates occurs upstream of mitotic CDK phosphorylation, or that Aurora kinases play an under-recognised role during DNA replication. Indeed, it has previously been suggested that Aurora kinases regulate DNA synthesis through their activity in mitosis, by chromatin remodeling and by organising the replication origin firing program (Koch et al., 2011). Further studies will be required to elucidate interphase roles of Aurora kinases.

A fourth important finding is that, even though we confirm that phosphorylation in general tends to locate to IDRs (Iakoucheva et al., 2004), proteins undergoing cell cycle-regulated phosphorylation are more intrinsically disordered than all other phosphoproteins. This trend is even higher in *bona fide* CDK substrates in both human and yeast. Interestingly, a recent large-scale proteogenomics study also found that cell cycle-dependent proteins are enriched in disorder, while revealing that most are regulated at the post-translational, rather than transcriptional, level (Mahdessian et al., 2021). Along with our results, this suggests that cell cycle regulatory mechanisms have been selected to control intrinsically disordered proteins.

This could be related to cell cycle control of MLOs, many of which break down in mitosis. Since CDK sites tend to cluster together in IDRs (Holt et al., 2009; Moses et al., 2007), and our data shows that many cell cycle-regulated CDK substrates are key components of MLOs, it is likely that CDK-mediated phosphorylation will affect MLO structure. Indeed, CDKs can control formation of stress granules (Yahya et al., 2021) and histone bodies (Hur et al., 2020), and possibly other MLOs. An emerging model is that liquid-liquid phase separation, which depends on weak interactions between IDRs, underlies the self-assembly of many MLOs (Banani et al., 2017; Hyman et al., 2014; Woodruff et al., 2018). Thus, switch-like phosphorylation of some IDRs may promote rapid MLO disassembly in mitosis, acting analogously to a detergent that dissolves liquid phase boundaries. In support of this model, DYRK3 is a CDK-related kinase whose inhibition disrupts mitotic remodeling of stress granules, splicing speckles and pericentriolar material, all of which are thought to assemble via phase separation (Rai et al., 2018). It is not currently feasible to study effects of CDK-inhibition on mitotic MLO phosphorylation and structure since CDK inhibition prevents or reverses entry into mitosis. Nevertheless, the data generated here pave the way for future *in vitro* studies on the effects of CDK-mediated phosphorylation on potential phase separation of key IDPs of MLOs.

In conclusion, this work reveals *in vivo* that cell cycle dynamic and CDK sites are both quantitatively and qualitatively different from other phosphosites: they are the most numerous, they occur in a switch-like manner at mitosis on multiple proteins of complexes involved in diverse cellular processes, and they are highly enriched within IDRs. Overall, our data suggest that evolution selected IDRs to allow coordination of diverse cellular processes during the cell cycle, and also chose CDKs as principal regulators of their phosphorylation state. In this view, cell cycle control may be less specific than previously assumed.

## Acknowledgments

We thank Merlijn Witte for technical assistance with the *Xenopus laevis* egg fertilization experiments, Ariane Abrieu for a gift of CSF egg extracts, and Markus Raschle from the Technical University of Kaiserslautern for providing the *Xenopus laevis* protein database.

## Funding

AJRH and MA acknowledge support from the Horizon 2020 program INFRAIA project Epic-XS (Project 823839) and the NWO funded Netherlands Proteomics Centre through the National Road Map for Large-scale Infrastructures program X-Omics (Project 184.034.019) of the Netherlands Proteomics Centre. JMV is supported by scholarships from the Ministry of Science and Technology of Costa Rica (MICITT) and the University of Costa Rica (UCR). PK and MV are funded by the Oncode Institute which is financed by the Dutch Cancer Society and by the gravitation program CancerGenomiCs.nl from the Netherlands Organisation for Scientific Research (NWO). DF and LK are Inserm employees. GD is funded by the Institut National de Cancer, France (INCa) PRT-K programme (PRT-K17 n° 2018-023). The Fisher lab is funded by the Ligue Nationale Contre le Cancer, France (EL2018.LNCC/DF) and INCa (PLBIO18-094).

## Author contributions

MA and DF conceived and supervised the project. JMV, PK and LK designed and interpreted experiments. JMV, HT, AH, GvM and GD performed experiments and interpreted the data. MV supervised GvM. JMV, LK, GD, DF and MA wrote the paper.

## Competing interests

Authors declare no competing interests.

## Data and materials availability

All data is available in the main text or the supplementary materials. The mass spectrometry data have been deposited via the PRIDE partner repositories. The proteome data to the ProteomeXchange Consortium (PXD026076) and the targeted proteomics data to Panorama Public (PXD026088). All other data, code, and materials are available on request.

Reviewer login:

ProteomeXchange; Username, Password: Ya2mrP0H

Panorama Public; Username, Password: IJSwZMnM

## Supplementary Materials

Materials and Methods

Figures S1-S6

Data S1-S4

## Notes

### Competing Interest Statement

The authors have declared no competing interest.

