## Supplemental Information for "Single-embryo phosphoproteomics reveals the importance of intrinsic disorder in cell cycle dynamics"

**Affiliations:**

†‡ Equal contributions

**Supplementary Materials:**

Figures S1-S6

Materials and Methods

Data S1 - Phosphoproteomics-data
Data S2 - CDK-phosphorylation-data
Data S3 - Protein-disorder-prediction
Data S4 - Human-proteins-in-MLOs

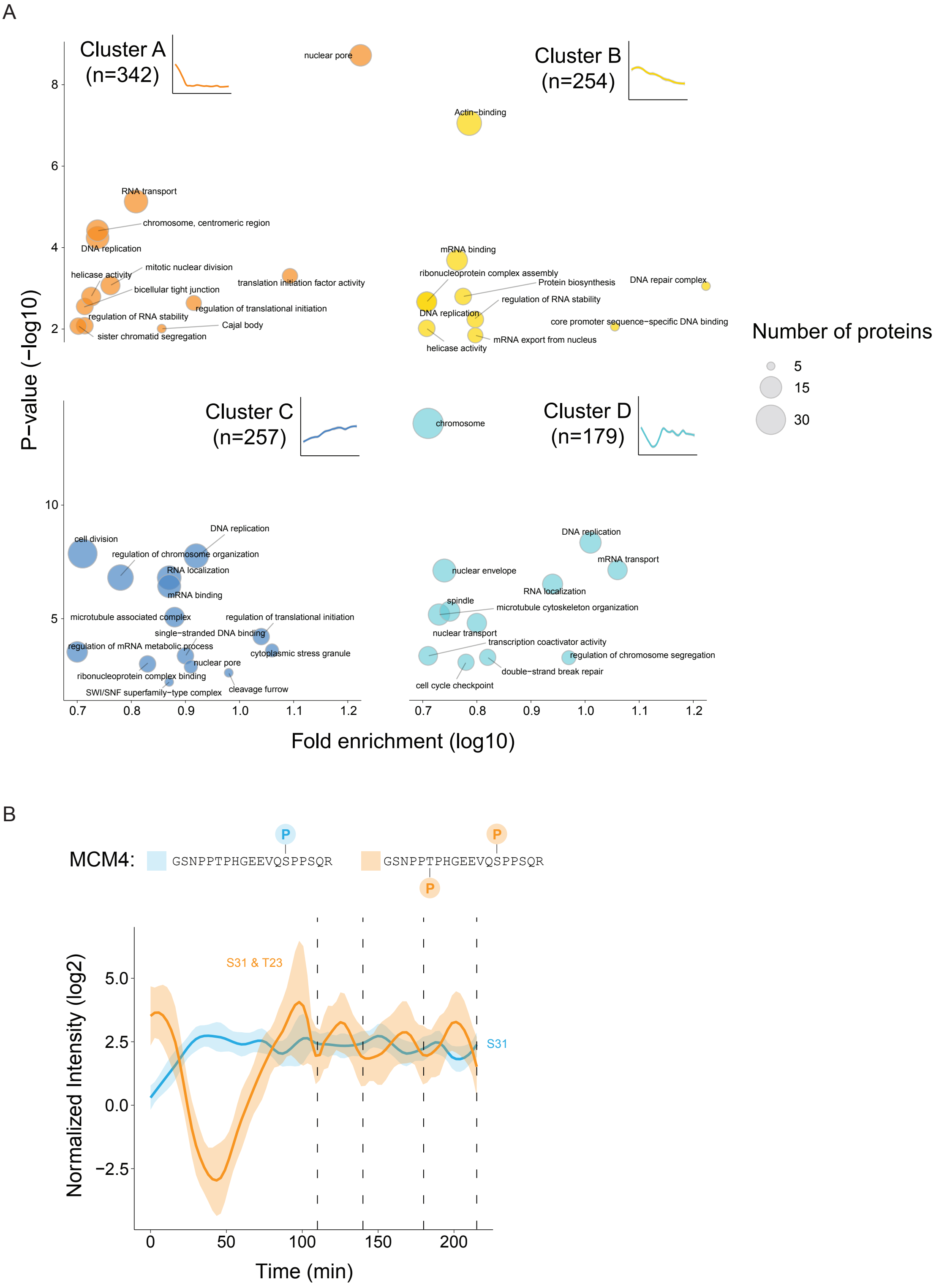


**Figure S1. Phosphosite dynamics corelates with cell cycle phases, related to Figure 1.** (A) Scatter plots of significantly enriched (Fisher's exact test with Bonferroni correction, p<0.05) GO (BP, MF, CC, Uniprot keywords) terms for all dynamic phosphosites per cluster in the *in vivo* experiment, presenting the fold-enrichment of specific terms vs statistical significance. The size of the circles correlates with the number of proteins associated with the specific term. More details and the full list of enriched GO terms per cluster is found in Data S1. (B) *In vivo* reciprocal trends of singly- and multi-phosphorylated peptides carrying phosphorylated T23 and S31 of MCM4 (dashed lines depict the time points of cell division): orange curves, the trend of T23 and S31 in the multi-phosphorylated peptide; blue curve, the trend of S31 in the singly-phosphorylated peptide.

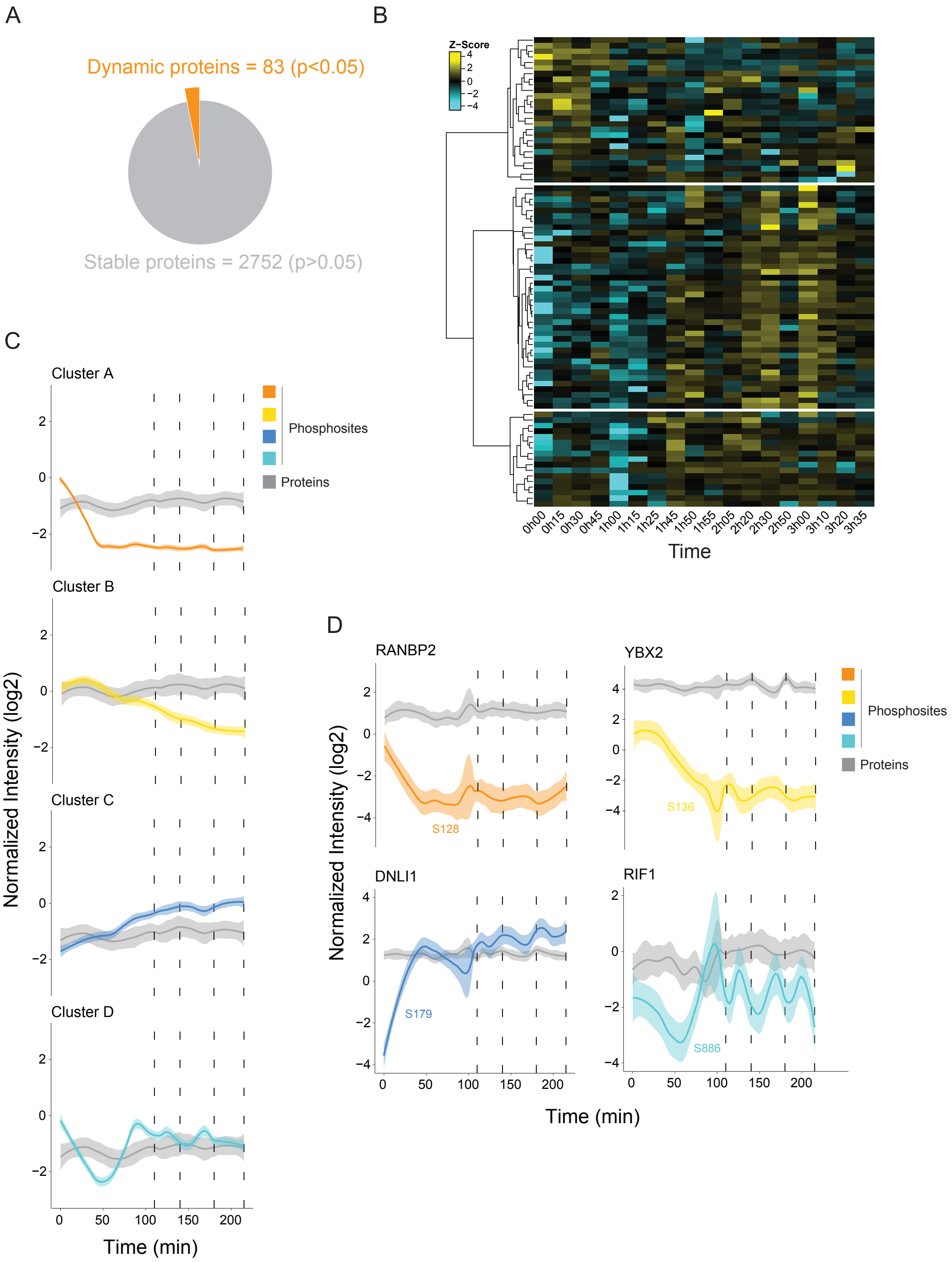

**Figure S2. Variation in the total proteome versus the phosphoproteome in early embryonic cell cycles, related to Figure 1.** (A) Total proteome analysis reveals 83 proteins out of 2835 showing significant changes in abundance (ANOVA, Benjamini-Hochberg correction, FDR 0.05) over the time course. (B) Heat map showing abundance of the variable proteins over the time course. (C) Comparison of dynamic variations in total protein compared to total phosphosites from the four clusters shown in Figure 1 (dashed lines depict the time points of cell division). (D) Examples of dynamics of individual phosphosites from the four clusters shown in Figure 1 and levels of the corresponding protein (dashed lines depict the time points of cell division).

A

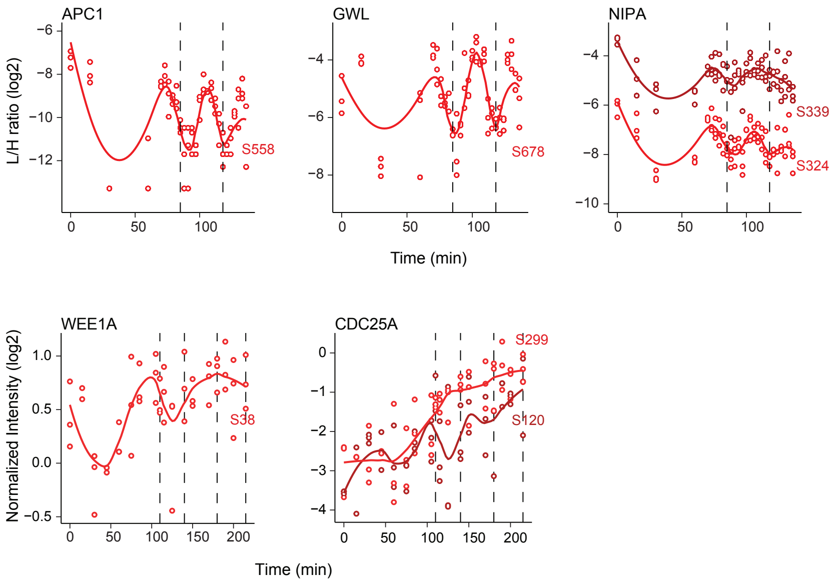


B

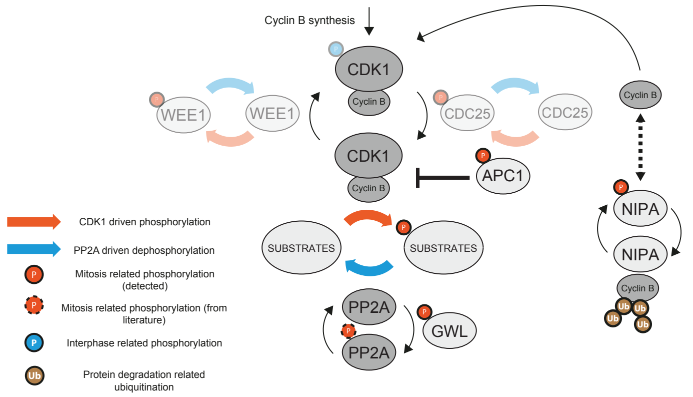


**Figure S3. Phosphorylation dynamics of the CDK1-oscillator network, related to Figure 2.** (A) Single phosphosite plots of CDK1 regulators measured by targeted (top) or shotgun (below) phosphoproteomics. (B) CDK1-oscillator network: our data suggests that control of cyclin levels via positive (*e.g.* NIPA ubiquitin ligase) and negative (*e.g.* APC) feedback loops, accompanied by PP2A inactivation via GWL, can generate oscillation of CDK1 activity during early cell divisions. CDK1 Y15 regulation via feedback loops consisting of CDC25 and WEE1A (greyed out) seems to be less important for switch-like mitotic phosphorylation after the first cell division.

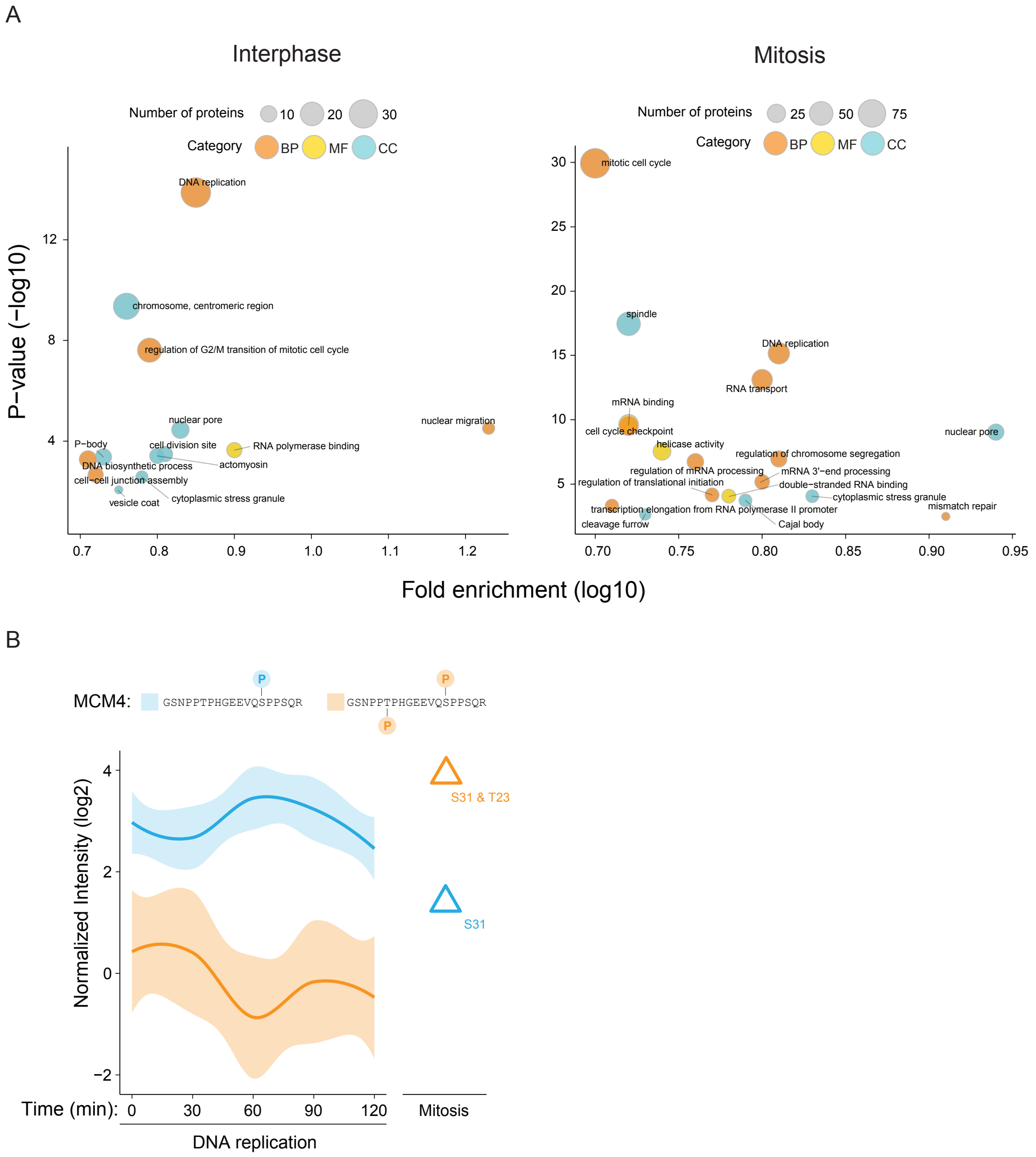


**Figure S4. *In vitro* phosphoproteomics discriminates interphase and mitotic phosphorylation, related to Figure 3.** (A) Scatter plots of significantly enriched (Fisher's exact test with Bonferroni correction, $p<0.05$) GO terms for all dynamic phosphosites upregulated during interphase (left) and mitosis (right), presented as fold-enrichment of specific terms vs statistical significance. The size of the circles correlates with the number of proteins associated with the specific term, while the color corresponds to the GO term category. (B) *In vitro* dynamics of T23 and S31 of MCM4. Orange curve shows upregulation of the multi-phosphorylated peptide (T23 and S31) during mitosis, while the blue curve shows the opposite trend for the singly-phosphorylated peptide (S31), as observed *in vivo*, Figure S1.

A

| | PLK | AURKA/B | CK1/2 | DDK | CDK full | CDK min |
|---|---|---|---|---|---|---|
| Cluster A (n=342) | 14 | 12 | 97 | 12 | 14 | 205 |
| Cluster B (n=254) | 11 | 27 | 67 | 12 | 16 | 124 |
| Cluster C (n=257) | 9 | 29 | 72 | 31 | 36 | 135 |
| Cluster D (n=179) | 7 | 5 | 51 | 4 | 31 | 150 |
| Total (n=1032) | 41 | 73 | 287 | 59 | 97 | 614 |

| | | PLK | AURKA/B | CK1/2 | DDK | CDK full | CDK min |
|---|---|---|---|---|---|---|---|
| M-phase | Cluster 1 (n=161) | 23 | 7 | 56 | 5 | 12 | 67 |
| | Cluster 2 (n=172) | 5 | 6 | 62 | 6 | 23 | 114 |
| | Cluster 3 (n=702) | 16 | 23 | 224 | 5 | 123 | 565 |
| Interphase | Cluster 4 (n=151) | 13 | 52 | 48 | 8 | 6 | 20 |
| | Cluster 5 (n=433) | 15 | 34 | 136 | 67 | 25 | 196 |
| | Cluster 6 (n=109) | 6 | 9 | 36 | 14 | 9 | 40 |
| | Total (n=1728) | 78 | 131 | 562 | 105 | 198 | 1002 |

B

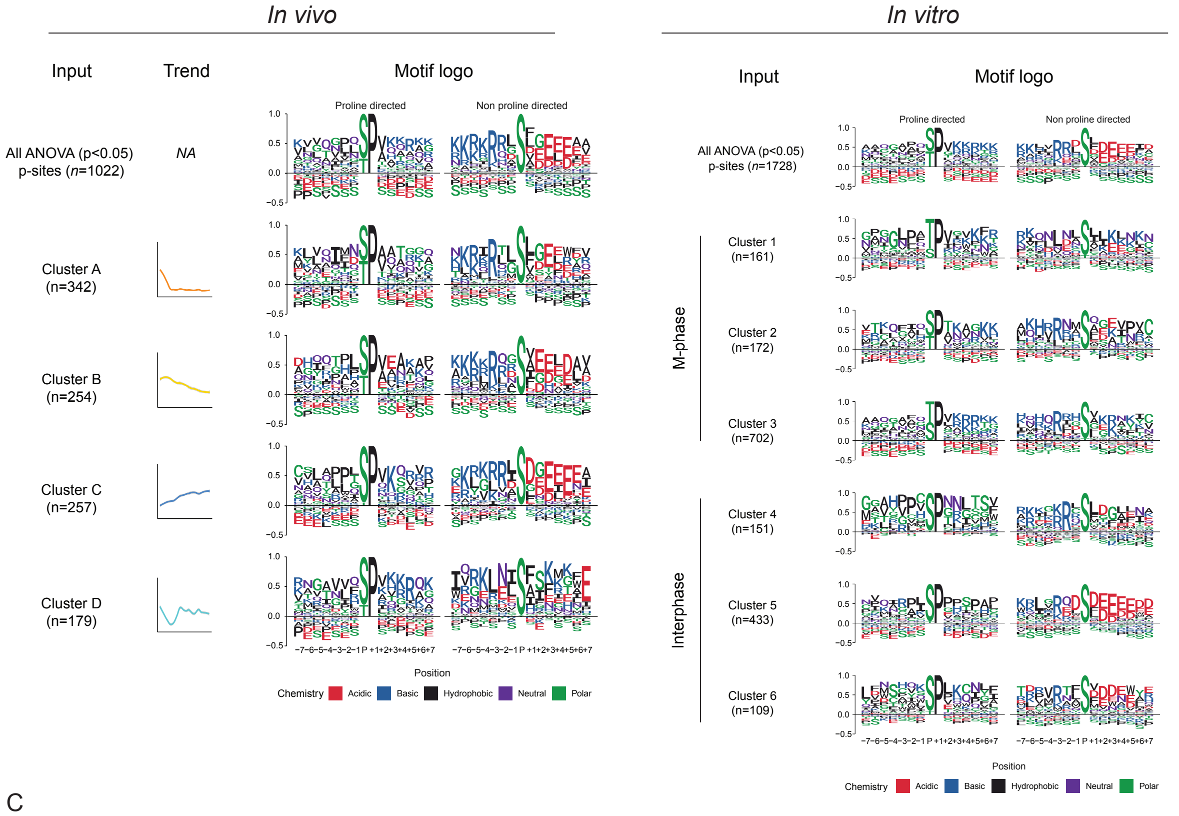


C

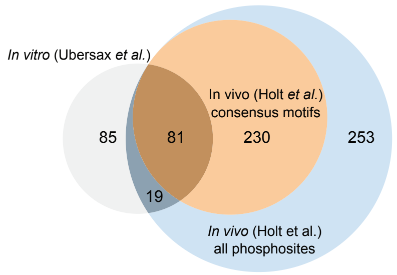


**Figure S5. Phosphorylation motif analysis, related to Figures 4 and 5.** (A) Observed phosphorylation motifs in the dynamic phosphoproteome *in vivo* (left) and *in vitro* (right). See methods for details. Note: in some cases, the sum of consensus sites exceeds the number of phosphosites due to redundancy between motif predictions. (B) Sequence motif logo for all dynamic phosphosites and for each of the clusters shown in Figure 1 and Figure 3, for the *in vivo* and *in vitro* experiments, respectively. Motifs are shown separately for proline-directed and non-proline directed phosphosites. (C) Venn diagram of observed *in vitro* (grey) and *in vivo* (blue) yeast CDK targets. *In vivo* targets showing CDK minimal consensus motif phosphorylations are highlighted in orange.

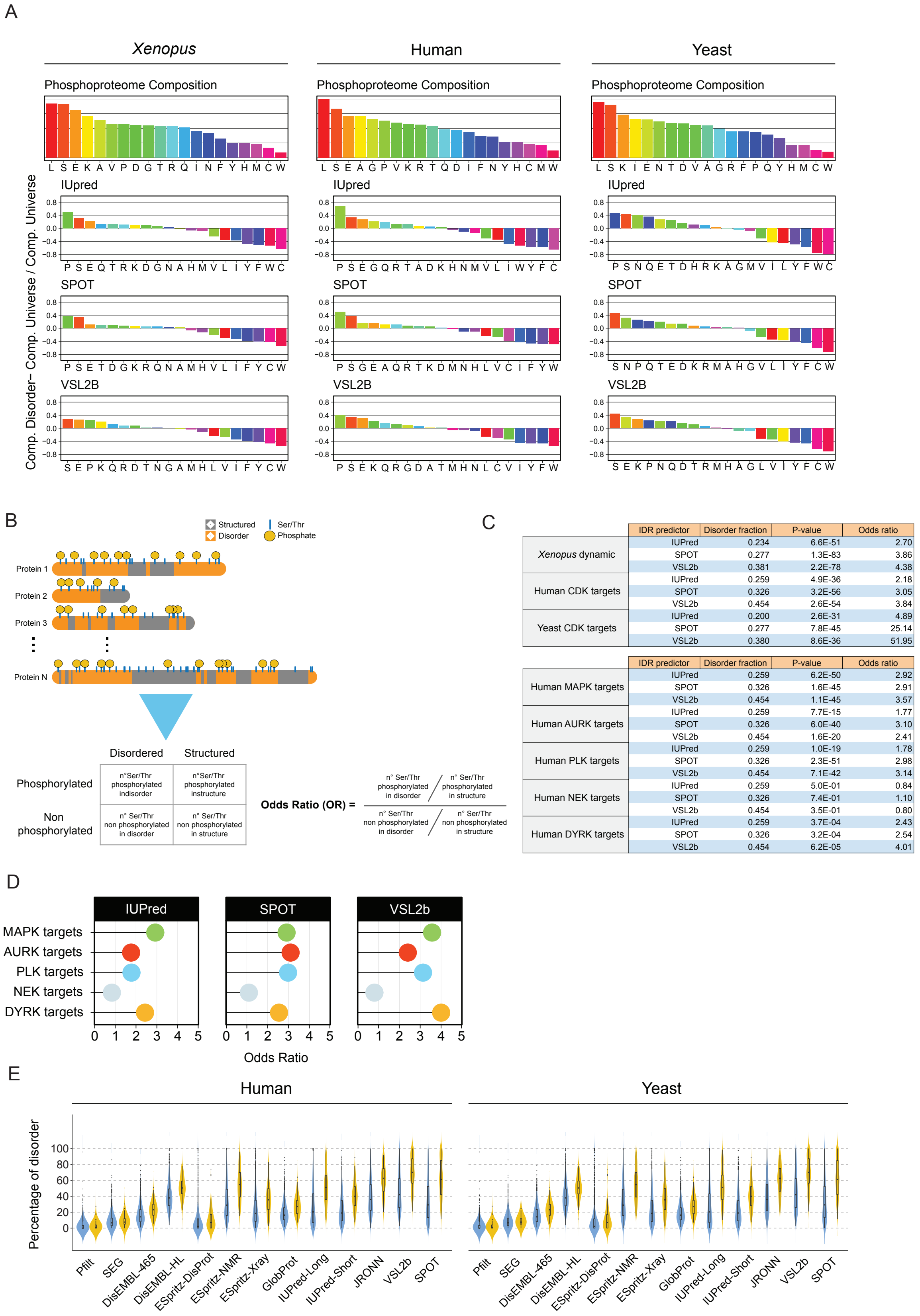


| | IDR predictor | Disorder fraction | P-value | Odds ratio |
|---|---|---|---|---|
| *Xenopus* dynamic | IUPred | 0.234 | 6.6E-51 | 2.70 |
| | SPOT | 0.277 | 1.3E-83 | 3.86 |
| | VSL2b | 0.381 | 2.2E-78 | 4.38 |
| Human CDK targets | IUPred | 0.259 | 4.9E-36 | 2.18 |
| | SPOT | 0.326 | 3.2E-56 | 3.05 |
| | VSL2b | 0.454 | 2.6E-54 | 3.84 |
| Yeast CDK targets | IUPred | 0.200 | 2.6E-31 | 4.89 |
| | SPOT | 0.277 | 7.8E-45 | 25.14 |
| | VSL2b | 0.380 | 8.6E-36 | 51.95 |

| | IDR predictor | Disorder fraction | P-value | Odds ratio |
|---|---|---|---|---|
| Human MAPK targets | IUPred | 0.259 | 6.2E-50 | 2.92 |
| | SPOT | 0.326 | 1.6E-45 | 2.91 |
| | VSL2b | 0.454 | 1.1E-45 | 3.57 |
| Human AURK targets | IUPred | 0.259 | 7.7E-15 | 1.77 |
| | SPOT | 0.326 | 6.0E-40 | 3.10 |
| | VSL2b | 0.454 | 1.6E-20 | 2.41 |
| Human PLK targets | IUPred | 0.259 | 1.0E-19 | 1.78 |
| | SPOT | 0.326 | 2.3E-51 | 2.98 |
| | VSL2b | 0.454 | 7.1E-42 | 3.14 |
| Human NEK targets | IUPred | 0.259 | 5.0E-01 | 0.84 |
| | SPOT | 0.326 | 7.4E-01 | 1.10 |
| | VSL2b | 0.454 | 3.5E-01 | 0.80 |
| Human DYRK targets | IUPred | 0.259 | 3.7E-04 | 2.43 |
| | SPOT | 0.326 | 3.2E-04 | 2.54 |
| | VSL2b | 0.454 | 6.2E-05 | 4.01 |

**Fig. S6. Cell cycle phosphorylation occurs predominantly in IDRs, related to Figure 5.** (A) Differential amino acid composition (see methods) in disordered regions for *Xenopus*, human and yeast determined with three IDR predictors. Amino acids are coloured in a rainbow pattern according to their relative abundance in each phosphoproteome. Disruptions of the rainbow pattern show specific compositional signatures for IDRs. (B) Scheme of the Odds Radio analysis using contingency tables. The counts of phosphorylated Ser/Thr for all the proteins for each set (CDK-mediated, yeast/human, or dynamic, *Xenopus*, and other cell cycle-related kinases in human), in disordered and structured regions, are stored in a 2x2 table. (C) Tables showing results of statistical analysis of Odds Ratio with Fisher's test, using three disorder predictors. (D) Plots of the Odds Ratio for human cell cycle-related non-CDK kinases and MAPK. (E) Violin plots of the distribution of percentage of disordered residues per protein for CDK targets *vs* the rest of the phosphoproteome for human and yeast. Intrinsic disorder information of 13 different predictors was obtained from MobiDB, except for SPOT (calculated).

### Materials and Methods

Egg collection and *in vitro* fertilisation

Female *X. laevis* frogs were primed with 50 international units (IU) of human chorionic gonadotropin at least 2 days, and no more than 7-8 days before a secondary injection with 625 IU to induce ovulation. Roughly 16 hours after the second injection, fresh eggs were collected by pelvic massage and kept in 1x Marc's Modified Ringer's (MMR). Next, eggs were placed in a petri dish and checked under the microscope to keep only those that exhibited the healthy pigment pattern (dark animal pole and white vegetal pole).

To perform the *in vitro* fertilisation, around 1/3 of a full testis was cut into fine pieces and mixed with in 500 µL of 1x MMR. The suspension was pipetted up and down until big clumps were dissolved. Next, buffer was removed from the petri dish and the eggs were collected. Once eggs were well dispersed across the dish, the sperm suspension was added. The dish was then flooded with 0.1x MMR to induce fertilisation.

Sample collection

The first time point was collected immediately before adding the sperm suspension (time 0' corresponds to the unfertilised egg). Eggs were kept at room temperature (18-20°) and under a dissecting microscope after fertilisation. At approximately 15 minutes, fertilised eggs underwent shrinkage of the animal hemisphere and rotation within the vitelline membrane, so that the animal hemisphere faced upwards. These changes are known indicators of successful fertilisation, so only the eggs that underwent these changes were used for the experiment.

Samples were collected approximately every 15 minutes. Eggs were rapidly placed in individual tubes and snap froze in liquid nitrogen, trying to preserve the phosphorylation events occurring at that specific time. Since the eggs were monitored under the microscope, we were able to determine if samples were collected before or after a cell division had occurred.

*Xenopus* egg extracts

Interphase *Xenopus* egg extracts were prepared, and DNA replication time courses performed, as described previously (Parisis et al., 2017). Mitosis was induced in extracts by adding recombinant GST-Cyclin BΔ90 (40 ng/ml).

Cell lysis

For the cell lysis, we used a similar approach to that described by Lindeboom *et al* (Lindeboom et al., 2018). Briefly, each sample was thawed and homogenised with 15µL of ice-cold cell lysis buffer (20mM Tris-HCl pH 8.0, 70mM KCl, 1mM EDTA, 10% glycerol, 5mM DTT, 0.125% Nonidet P-40, 1mM PMSF, 1xcomplete EDTA-free protease inhibitor, 1xPhoStop). Samples were subsequently centrifuged at max speed on a benchtop Eppendorf centrifuge. 10µL of soluble material was recovered and snap-frozen in liquid nitrogen. Samples were stored at -80°C until further processing.

Protein digestion and phosphopeptide enrichment

Cell lysates were digested using the FASP method (Wisniewski et al., 2009). Briefly: proteins were thawed and immediately reduced and alkylated with 10mM DTT and 0.05M iodoacetamide. Next, proteins were digested with Lys-C (overnight) at 37°C in a wet chamber, followed by addition of trypsin and further incubation under the same conditions for 4 hours. Both enzymes were used at 1:50 enzyme to protein ratio (protein quantification by Bradford assay showed that each individual egg provides ~20µg of yolk free protein). For the egg extract experiment, each FASP filter was loaded with 200µg of protein. Peptides were cleaned using the Oasis HLB 96 well plates (Waters Corporation) and consequently subjected to phosphopeptide enrichment using Fe(III)-NTA 5µL cartridges in the automated AssayMAP Bravo Platform (Agilent Technologies), as described by Post *et al* (Post et al., 2017). Both flow through (peptides) and eluates (phosphopeptides) were dried down and stored at -80°C until further use.

LC-MS/MS analysis

All samples for label-free shotgun proteomics were analysed using a UHPLC 1290 system (Agilent Technologies) coupled to an Orbitrap Q Exactive HF mass spectrometer (Thermo Fisher Scientific). Nano flow rate was achieved using a split flow setup aided by an external valve as described by Meiring *et al* (Meiring et al., 2002). Peptides were first trapped onto a pre-column (inner diameter [ID] of 100µm and 2cm length; packed in-house with 3µm C18 ReproSil particles [Dr. Maisch GmbH]) and eluted for separation into an analytical column (ID of 75µm and 50cm length; packed in-house with 2.7µm Poroshell EC-C18 particles (Agilent Technologies). The latter was done using a two-buffer system, consisting of buffer A (0.1% formic acid [FA] in water) and buffer B (0.1% FA in 80% ACN). Peptides were trapped during 5 minutes at 5µL/min flow-rate with solvent A. For the measurement of the full proteome, we used a 155 minute gradient from 10 to 36% of solvent B. For the phosphoproteome we used a 95 minute gradient from 8 to 32% of solvent B. Both methods included a wash with 100% solvent B for 5 minutes followed by a column equilibration with 100% solvent A for the last 10 minutes.

The mass spectrometer was operated in data dependent acquisition (DDA) mode. Full scan MS was acquired from 375-1600m/z with a 60,000 resolution at 200m/z. Accumulation target value was set to 3e6 ions with a maximum injection time of 20ms. Up to 15 (12 for the phosphoproteome) of the most intense precursor ions were isolated (1.4m/z window) for fragmentation using high energy collision induced dissociation (HCD) with a normalised collision energy of 27. For MS2 scans an accumulation target value of 1e5 ions and a maximum injection time of 50ms were selected. Scans were acquired from 200-2000m/z with a 30,000 resolution at 200m/z. Dynamic exclusion was set at 24s for the proteome and 12s for the phosphoproteome.

For targeted proteomics, an EASY-nLC 1200 System (Thermo Fisher Scientific) coupled to an Orbitrap Q Exactive HF was used. Peptides were separated using an EASY-Spray analytical column (ID of 75µm and 25cm length; packed with 2µm C18 particles with a 100 Å pore size) (Thermo Fisher Scientific). Gradient lengths were shortened to 60 minutes. Phosphopeptides of interest from the previous experiment were selected and heavy-labeled versions were synthesised (JPT Peptide Technologies). These synthetic standards were used during method development for retention time scheduling and instrument ion fill-time optimisation. Additionally, synthetic heavy peptides were pooled and combined with synthetic retention time peptide standards (iRT, Biognosys) to generate a spectral library, measured in DDA mode using the same LC-MS setup. This spectral library provided fragment intensity and retention time information for quality control assessment of targeted measurements. Samples were reconstituted in 2% FA containing ~200fmol of each synthetic standard. The mass spectrometer was operated in data independent acquisition mode with an inclusion list of targets for parallel reaction monitoring (PRM). The list included the m/z values for the heavy and light versions of the phosphopeptides. Optimal measurement parameters were determined using test samples spiked with the heavy-labeled standards in order to guarantee optimal sensitivity for detection of endogenous phosphopeptides. We measured the targets of interest in a scheduled fashion, during a four-minute window with a 120,000 resolution, maximum injection time of 246ms and an accumulation target value of 2e5 ions, to ensure maximum specificity and sensitivity.

Data processing

DDA raw files were processed with MaxQuant (Cox and Mann, 2008) (v1.6.0.1) using a false discovery rate (FDR) <0.01. The default settings were used, with the following exceptions: variable modifications, specifically methionine oxidation, protein N-term acetylation and serine, threonine and tyrosine phosphorylation were selected. Cysteine carbamidomethylation was selected as a fixed modification. We also enabled the 'match between runs' option with the default values. Fractions were set so that matching was done only among biological replicates and samples of consecutive time points. The database search was conducted against a database generated by Temu *et al* (Temu et al., 2016). This was particularly helpful since other publicly available databases contained several incomplete and/or poorly annotated sequences, which proved to be impractical for further data analysis.

The data was uploaded to the Perseus platform (Tyanova et al., 2016) for further analysis. Briefly: decoy sequences and potential contaminants were filtered out. Only high confidence localisation (>0.75 localisation probability) phosphosites were conserved for further analysis. Intensities were log2 transformed and then normalised by subtracting the median intensity of each sample. Biological replicates were grouped accordingly by time point; this grouping allowed us to filter the data and keep only those phosphosites that could be detected in at least two out of three biological replicates in any of the time points. Missing values were imputed from a random normal distribution applying a downshift of 1.8 times the standard deviation of the dataset, and a width of 0.3 times the standard deviation. This effectively replaced missing values at the lower end of the intensity distribution. We then performed an ANOVA (Benjamini-Hochberg FDR <0.05) to determine which phosphosites displayed statistically significant changes through the time course. Average phosphosite intensities were grouped using a combination of k-means and hierarchical clustering using the ComplexHeatmap package (Gu et al., 2016) in R. Protein intensities were processed in a similar fashion, removing proteins that were only identified by peptides that carry one or more modified amino acids.

Next, the full list of proteins with significantly changing phosphosites were matched against the human Uniprot database using the Basic Local Alignment Search Tool (BLAST), to render Uniprot identifiers that were compatible with different Gene Ontology (GO) analysis tools. We used the STRING web tool (Szklarczyk et al., 2019) to gain insight into the relation amongst dynamically phosphorylated proteins. The full list of phosphoproteins was uploaded and analysed using default settings. Next, the protein network was loaded into Cytoscape for clustering and visualisation. Proteins were clustered using GLay community clustering (Su et al., 2010) and enrichment of GO terms per cluster was obtained using BiNGO (Maere et al., 2005) (shown in Fig. 1E).

GO term enrichment was also acquired individually for each group (A-D) obtained after hierarchical clustering of dynamic phosphosites. For this we used STRING and filtered the enriched GO terms to keep only those with p <0.01, fold enrichment >5 and a minimum of 5 proteins per term. The list of terms was further condensed by removal of redundant terms using the Revigo web tool (Supek et al., 2011). Remaining GO terms (including BP, MF and CC) were manually curated to further avoid redundancy. The final set of GO terms per cluster are shown in supplementary figure 1. Following the same strategy, we analysed GO term enrichment for the interphase and mitotic clusters from the *in vitro* dataset separately (shown in Figure S4).

PRM raw files were analysed with Skyline software (MacLean et al., 2010). Signal quality for each target of interest was assessed visually for all samples. Quality control of endogenous signals was done by confirming the perfect co-elution of both peptide forms (heavy and light), assessing their retention time and peak shape. We also used the similarity of the relative intensity of fragment ions (rdotp > 0.9) between light and heavy to exclude signals that showed poor correlation. Quantifications were done with a minimum of three fragments per phosphopeptide. The data was loaded into R for data cleanup and visualisation, using the Complex-Heatmap and ggplot2 (Wickham et al., 2019) packages.

<u>Motif analysis</u>

Obtained phosphopeptides were aligned by centering them around the phosphosite detected and the conserved motifs for the different kinases were determined using regular expresions by applying  the following rules:

PLK: [D/N/E/Y]-X-[S/T]-[ Hydrophobic / ^P]

AURA/AURB: [K/R]-X-[S/T]*[^P]

NEK: [LIMIFIW]-X-[SIT]*-[AIVIIIILIFIWIYIM]-[KIR]

Casein kinase 1: [D/E]-[D/E]-[D/E]-X-X-[S/T]* or [S/T]-X-X-[S/T]*

Casein kinase 2:[S/T]-[S/T]*-X-[E/D/S]

DDK: [S/T]*-[E/D]-X-[E/D] or [S/T]*-[S/T]-P

PKA: R-[R/K]-X-[S/T]*-[Hydrophobic]

Cdk full consensus motif: [S/T]*-P-X-[K/R]

Cdk minimal consensus motif: [S/T]*-P

Where [] groups multiple amino acids for one position, ^ at the left of a certain amino acid informs that it is forbidden for that position, and X represents any amino acid.

<u>Data collection for human and yeast CDK1 targets.</u>

Data of CDK1 substrates for *S. cerevisiae* were downloaded from online supplementary information of papers describing two different studies using *in vitro* (4) and *in vivo* (26) approaches, respectively. We defined high confidence yeast CDK1 targets as the intersection of both datasets. Other phosphorylations detected in both studies for which there was no evidence for CDK1 involvement were considered as the non-CDK1-mediated phosphoproteome (universe). For human CDK1 subfamily targets, we extracted information available in the PhosphoSitePlus database (Hornbeck et al., 2015). An additional step of manual curation from the following studies (Bartsch et al., 1999; Blethrow et al., 2008; Chi et al., 2008, 2020; Curtis et al., 2002; Fourest-Lieuvin et al., 2006; Goto et al., 2006; Hégarat et al., 2020; Kitzmann et al., 1999; Klein et al., 2008; Li et al., 2006; Linder et al., 2017; Liu et al., 2000; Lowe et al., 1998; Milner et al., 1993; Orthwein et al., 2014; Thiel et al., 2002; Wyatt et al., 2013; Yun et al., 2003) was performed to obtain a high confidence human CDK1 subfamily targets dataset. The phosphoproteome universe was constructed with all the phosphorylated proteins deposited in the PhosphoSitePlus database after subtraction of the CDK1 subfamily targets.

Data is available in Data S2.

Data collection for MLO proteomes

Data from proteomics studies of the composition of MLOs characterised by liquid-liquid phase separation was obtained from the following sources: stress granules (Fong et al., 2013; Jain et al., 2016), nuclear speckles (Dopie et al., 2020; Fong et al., 2013), PML nuclear bodies (Fong et al., 2013; Liu et al., 2010), P-bodies (Hubstenberger et al., 2017), nucleoli (Stenström et al., 2020; Tafforeau et al., 2013), nuclear pore complexes (Lin and Hoelz, 2019), Cajal bodies (Fong et al., 2013; Machyna et al., 2013), Super-enhancer-Mediator condensates (Quevedo et al., 2019). Data is available in Data S4.

Prediction of intrinsically disordered regions.

For the UniProt proteomes of human, yeast and *Xenopus laevis*, disorder information was fetched from MobiDB (Piovesan et al.) with the exception of SPOT disorder predictor, which was calculated for all the proteins of each dataset. For Xenopus proteomics studies, we used the available standalone software of IUPred, VSL2B, and SPOT to predict IDRs in all the proteins of the database. Data is available in the Data S3.

Differential disorder composition

For the three organisms analysed (*Xenopus,* human and yeast ), the amino acid composition for the entire phosphoproteome and for the disordered regions of the phosphoproteome was calculated. For each amino acid, we estimated the differential disorder composition with the equation:

(Comp. Disorder – Comp. Phosphoproteome) / Comp. Phosphoproteome

Positives values show aminoacids enriched in disordered regions while negative values represent aminoacids depleted in disordered regions.

Bioinformatic and statistical analysis of disorder and phosphorylation

All the statistical analysis was performed with the R programming language (https://www.r-project.org/) using R studio as an integrated development environment (available at https://rstudio.com/). The packages Tidyverse (Wickham et al., 2019) and Bioconductor (Gentleman et al., 2004) were used for cleaning, manipulation, and graphical representation of the data. Sequence logos were generated using the information content as described (Douglass et al., 2012). IUPred scores were plotted with an *ad hoc* designed script, available upon request.

For the contingency table analysis, the disordered regions of CDK targets and dynamic phosphoproteins were calculated with three predictors (IUPred, VSL2B, and SPOT). For each combination of disorder predictor and phosphorylation dataset, a two-by-two table with the counts of phosphorylatable residues (Ser/Thr) phosphorylated or not, either located in IDRs or in structured regions (Fig. S6B), was generated. Each table was then analysed with the Fisher test for obtaining the odds ratio and the associated P-value.

The source code for all the analysis conducted in this publication is available upon request.

R: The R Project for Statistical Computing.

RStudio I Open source & professional software for data science teams.
